# Mammary ductal epithelium controls cold-induced adipocyte thermogenesis

**DOI:** 10.1101/2020.11.14.378687

**Authors:** Luis C. Santos, Douglas Arneson, Alexandra Alvarsson, Karthickeyan Chella Krishnan, Alessia Centzone, Sanil Patel, Shani Sadeh, In Sook Ahn, Graciel Diamante, Ingrid Cely, Atul J. Butte, Cédric Blanpain, Sarah A. Stanley, Aldons J. Lusis, Xia Yang, Prashant Rajbhandari

## Abstract

Sympathetic activation during cold exposure increases adipocyte thermogenesis via expression of mitochondrial protein uncoupling protein 1 (UCP1)^1^. The propensity of adipocytes to express UCP1 is under a critical influence of the adipose microenvironment and varies among various fat depots^2–7^. Here we report that cold-induced adipocyte UCP1 expression in female mouse subcutaneous white adipose tissue (scWAT) is regulated by mammary gland ductal epithelial cells in the adipose niche. Single cell RNA-sequencing (scRNA-seq) show that under cold condition glandular alveolar and hormone-sensing luminal epithelium subtypes express transcripts that encode secretory factors involved in regulating adipocyte UCP1 expression. We term mammary duct secretory factors as “mammokines”. Using whole-tissue immunofluorescence 3D visualization, we reveal previously undescribed sympathetic nerve-ductal points of contact and show that sympathetic nerve-activated mammary ducts limit adipocyte UCP1 expression via cold-induced mammokine production. Both *in vivo* and *ex vivo* ablation of mammary ductal epithelium enhances cold-induced scWAT adipocyte thermogenic gene program. The mammary duct network extends throughout most scWATs in female mice, which under cold exposure show markedly less UCP1 expression, fat oxidation, energy expenditure, and subcutaneous fat mass loss compared to male mice. These results show a previously uncharacterized role of sympathetic nerve-activated glandular epithelium in adipocyte thermogenesis. Overall, our findings suggest an evolutionary role of mammary duct luminal cells in defending glandular adiposity during cold exposure, highlight mammary gland epithelium as a highly active metabolic cell type, and implicate a broader role of mammokines in mammary gland physiology and systemic metabolism.

## MAIN

The scWAT depots in female mice are mostly mammary gland WATs (mgWAT) which is highly heterogenous tissue consisting of adipocytes, preadipocytes, mesenchymal stem cells, immune cells, endothelial cells, SNS nerve fibers, and mammary epithelial cells forming a ductal structure. The epithelial cells are divided into myoepithelial/basal cells, and luminal cells, which can be luminal hormone sensing, or not^8^. In virgin female mice, the mammary gland already has ductal structures in the anterior and posterior scWAT and metabolic cooperativity between luminal ductal cells and stroma is known to be important for mammary gland function and development^8,9^. Profound changes in mammary ducts and adipocytes are seen during gestation, pregnancy, lactation, and post-involution^10,11^. The importance of adipocytes for mammary duct morphogenesis, the dedifferentiation of adipocytes during lactation, and reappearance during lactation post-involution, all suggest a dynamic homeostatic interplay between ductal luminal epithelial cells and adipocytes^11–13^. It is not clear, however, what paracrine signaling programs from mammary ducts regulate adipocyte metabolism and thermogenesis. Notably, our current-state-of knowledge of WAT thermogenesis and UCP1 expression is mostly based on male scWATs, which lacks mammary glandular epithelial cells. Importantly, besides the role of immune cells in the adipose microenvironment, the contribution of other cell types in controlling adipocyte UCP1 expression is still not clear.

To study cellular heterogeneity, inter-tissue communication, and cellular transcription dynamics in mgWAT in a thermogenic condition, we isolated the stromal vascular fraction (SVF) from the mgWAT of 10-week-old virgin female mice exposed to 24-hour cold (COLD, 4°C) or room temperature (RT) and performed scRNA-seq (Fig. 1A). We obtained 12,222 cells and used Cell Ranger software from 10X Genomics for data processing and the R package Seurat^14^ to generate cell clusters and resolve their identities as previously described^15^ (see Methods). We integrated our dataset with eight other publicly available single cell datasets, from mammary gland tissues including *Tabula Muris* and *Tabula Muris Senis*^16–19^: i) for cell type identification, ii) to increase confidence in the projected cell type, iii) for sex and age differences, and iv) for mammary gland development (Extended Data Fig. 1A, and Table 1). This integrated dataset allowed us to precisely annotate various cell types present in mgWAT of female mice. Further subclustering of our integrated dataset based on known cell type marker genes identified clusters of a) adipocyte precursor cells (APCs), b) B cells, c) macrophages, d) T cells, e) endothelial cells, f) immune precursor cells (IPC), g) dendritic cells (DC), h) Schwann cells, i) myoepithelial cells (myoep), and j) luminal-hormone sensing (Luminal-HS), luminal-alveolar (Luminal-AV), luminal-HS-AV, and myoepithelial cells (Fig. 1B, Extended Data Fig. 1B and 1C).

**Figure 1.**
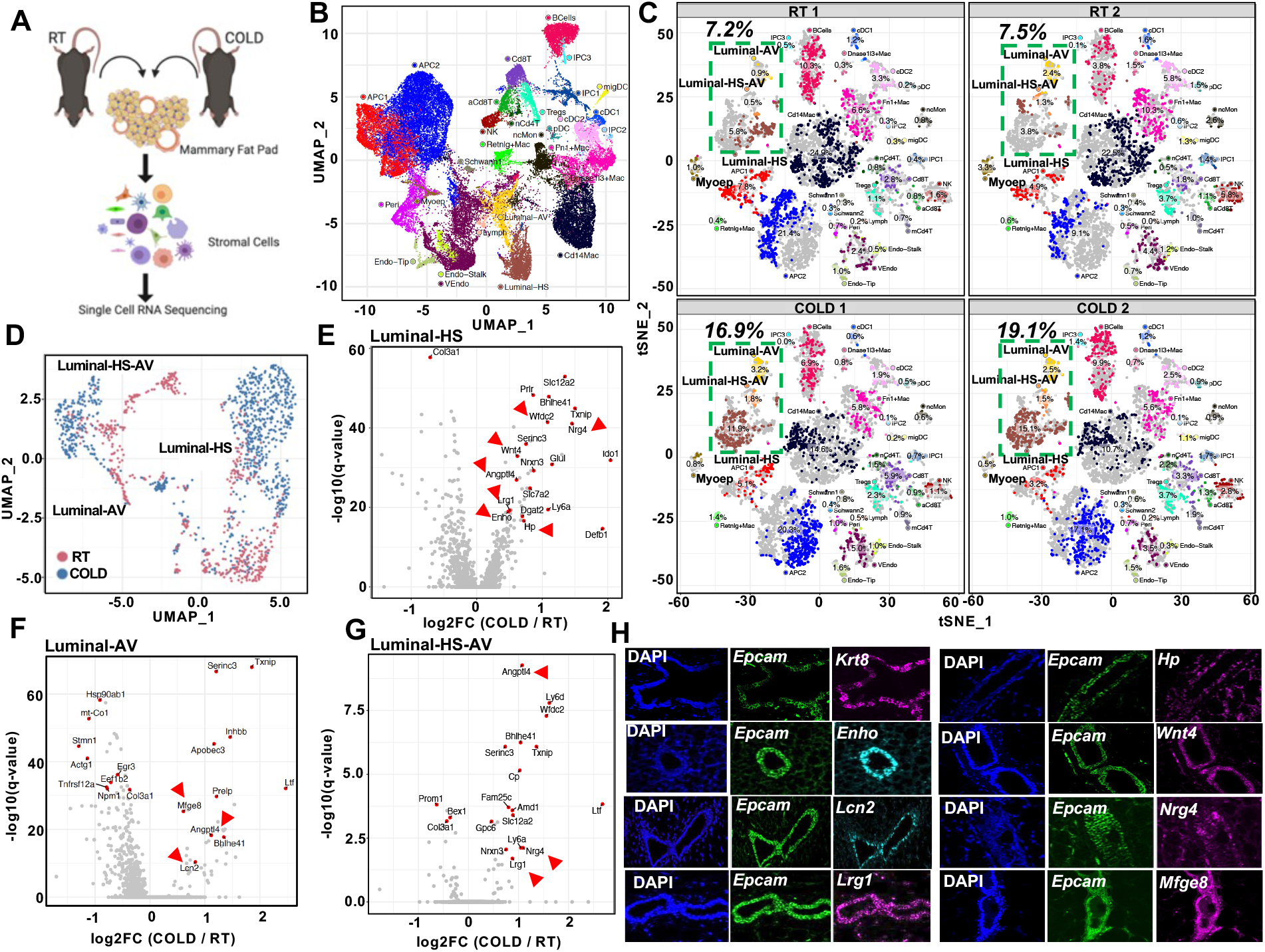
Deconstruction of mgWAT shows cold-induced remodeling of mammary epithelium. **(A)** Cartoon depiction of the scRNA-seq workflow showing isolation of stromal vascular fraction (stromal cells) from mammary fat pad (mgWAT) of 10-week-old 24 h RT or COLD exposed female mice. **(B)** UMAP plots of integrated single cell data from this study and 8 external datasets (see Methods). Each point represents a single cell and clusters are colored by cell type**. (C)** t-SNE plot of single cells from mammary gland and surrounding SVF colored by cell type and separated by sample. Relative fractions of each cell type in each sample are indicated on each cluster. Room temperature (RT) samples are on the top row and 4°C (COLD) are on the bottom row. **(D)** UMAP plot of luminal epithelial cell types from RT or cold mgWATs. **(E-G)** Differentially expressed genes between COLD treated mice and RT animals across Luminal-HS (E), Luminal-AV (F), and Luminal-HS-AV (G). Select significant DEGs (adjusted p-value < 0.05) are highlighted with the average log fold change between 4 degree and RT indicated on the y-axis. Genes indicated by red arrows encode for secreted factors. **(H)** RNAScope FISH (see Materials and methods) of indicated probes from mgWAT of 24 hr cold exposed mice. Abbreviations: APC, adipose precursor cells; IPC, immune precursor cells; Mac, macrophages; ncMon, non-classical monocytes; cDC, conventional dendritic cells; migDC, migratory dendritic cells; pDC, plasmacytoid dendritic cells; Tregs, regulatory T cells; mCd4T, memory Cd4 T cells; nCd4T, naïve Cd4 T cells; aCd8T, activated Cd8 T cells; Myoep, myoepithelial cells; VEndo, vascular endothelial cells; Endo-Tip, endothelial tip cells; Endo-Stalk, endothelial stalk cells; Lymph, lymphatic endothelial cells; Peri, pericytes; Luminal-HS, hormone-sensing luminal cells; Luminal-AV, secretory alveolar luminal cells; Luminal-HS-AV.

**Table 1:**
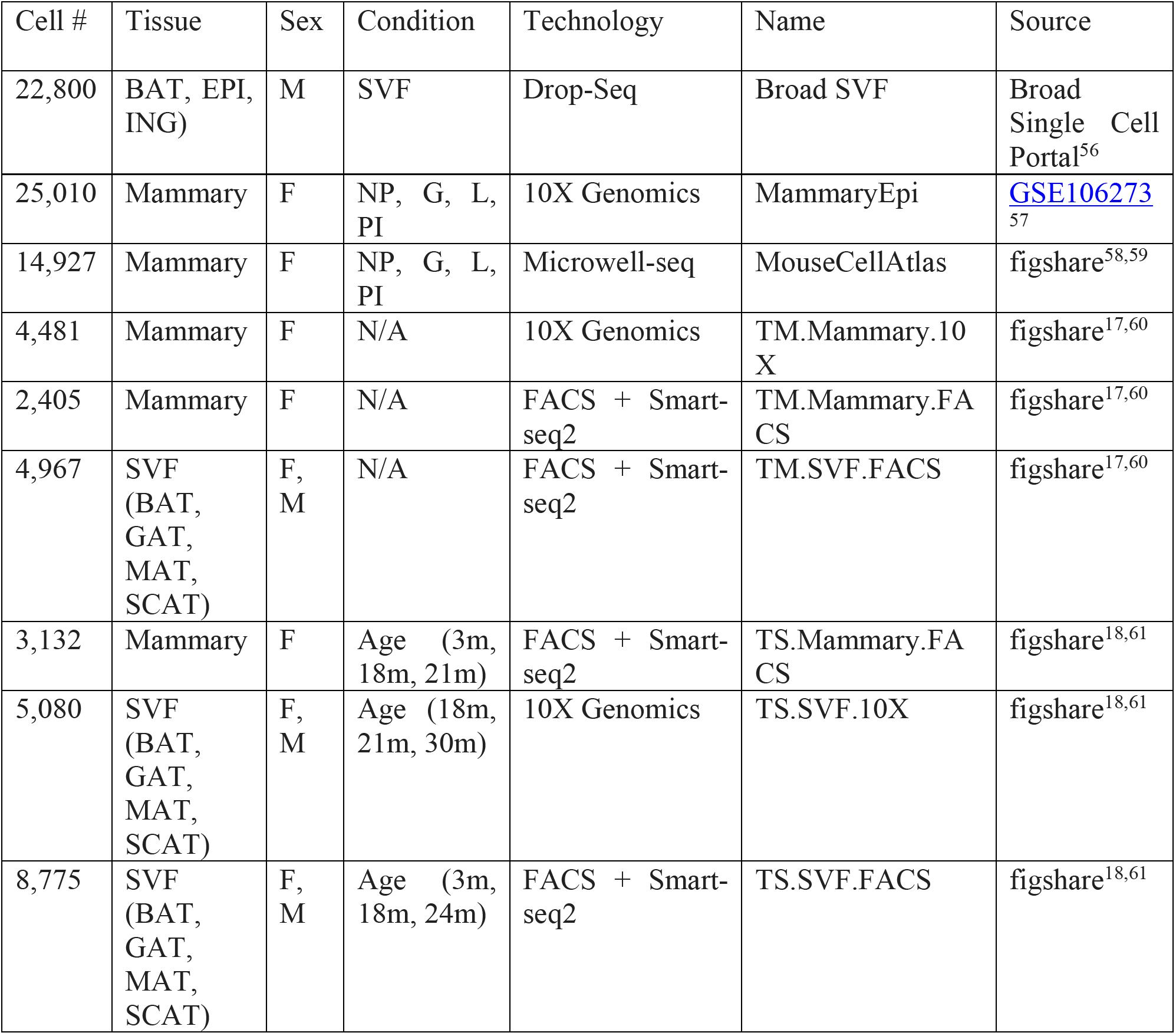
Publicly available single cell datasets used in this study. BAT – brown adipose tissue, EPI – epididymal white adipose tissue, ING – inguinal white adipose tissue, NP - nulliparous, G - gestation, L – lactation, PI - post involution, GAT – gonadal adipose tissue, MAT – mesenteric adipose tissue, SCAT – subcutaneous adipose tissue

To gain insight into the remodeling of stromal cells under adrenergic stress, we segregated the cumulative tSNE-plot into RT and COLD treatment by animal replicate. The tSNE and dot plots reveal global changes in relative proportions of SVF clusters between RT and COLD (Fig. 1C). Among all the clusters luminal cells showed significant differences in cell type percentages (RT 1:7.2%, RT 2: 7.5%; COLD 1: 16.9%, COLD 2: 19.1%) and appeared to have large differences in their global transcriptomic profiles in the t-SNE two-dimensional projection where cells from RT and COLD were segregated (Fig. 1C). To quantitatively determine the transcriptional impact of cold treatment on individual cell types, we characterized differentially expressed genes as a function of cluster types and found a high degree of transcriptional variation in luminal HS and AV under the cold conditions (Extended Data Fig. 1D). Further subclustering of luminal epithelial cell types (luminal HS, luminal AV, and luminal HS-AV) revealed marked differences in clustering at RT and COLD ^20^ (Fig. 1C and Extended Data Fig. 1D, 1E).

Luminal cell clusters showed remarkable transcriptional differences in cell clusters between RT and cold, implicating a potential remodeling of the luminal epithelium upon cold exposure (Fig. 1C, 1D and Extended Data Fig. 1D–1F). To probe for factors that are differentially expressed in luminal cells under cold exposure, we performed differential gene expression (DEG) analysis on RT and cold exposed luminal subclusters. We found upregulation of *Wnt4*, Adropin (*Enho*), leucine rich alpha-2 glycoprotein (*Lrg1*), Diglyceride acyltransferase (*Dgat2*), haptoglobin (*H*p), and angiopoietin-like 4 (*Angptl4*) in Luminal HS cells, lipocalin-2 (*Lcn2*), *Angptl4*, and Apolipoprotein B editing complex (*Apobec3*) in Luminal AV cells, and *Lrg1*, neuregulin 4 (*Nrg4*), ceruloplasmin (*Cp*), *Angptl4* in Luminal HS-AV cells (Fig. 1E–1G, Extended Data Fig. 1G). Many of these genes (shown by red arrows in Fig. 1E–1G) encode secreted factors that play important roles in local and systemic lipid metabolism^21–30^. t-SNE plots of normalized gene expression levels for cold-induced mammokines in mgWAT (our study), male scWAT SVFs and mature adipocytes^31^ show that *Angptl4* is also expressed by most mature adipocytes; however, other cold-induced mammokine genes showed relatively localized expression in ductal epithelial cell adhesion molecule (*Epcam*)+ cells (Extended Data Fig. 1G and 1H). Our RNAscope fluorescent in situ hybridization (FISH) analysis showed a highly localized expression of mammokines *Enho, Mfge8, Lrg1, Lcn2, Hp, Nrg4*, and *Wnt4* in *Epcam*+ and *Krt8*+ (luminal ductal epithelial markers) in mammary ductal luminal cells (Fig. 1H).

Shifts in the luminal epithelium transcriptomic state with cold and localized expression of beta-adrenergic receptors, *Adrb2* and *Adrb1* expression in luminal cells, suggests that these cells may directly respond to cold-induced SNS activation^32–34^ (Fig. 1C and Extended Data Fig. 1G). To determine if duct epithelial cells are innervated by sympathetic nerves, we used Adipoclear, a robust protocol based on immunolabeling-enabled three-dimensional imaging of solvent-cleared organ iDISCO (22), for high-resolution, three-dimensional imaging of mammary tissue. We analyzed the 3D distribution and density of a sympathetic marker, tyrosine hydroxylase (TH), and its relationship to EPCAM+ mammary ductal cells in mgWAT from mice exposed to RT or cold. Cold exposure caused prominent morphological changes in mammary ducts such as increased EPCAM+ branching and terminal ductal bifurcations (Fig. 2A and Extended Data Fig. 2A), which is consistent with data showing increased branch morphogenesis upon isoproterenol treatment^35^. Our data further reveal that nerve fibres are interwoven with mammary gland ducts and alveolar structures in mgWATs (Supplementary Movie 1). However, we did not see an effect of cold treatment on duct volume, nerve volume, or the ratio of duct-to-nerve volume (Fig. 2B). To examine interactions between sympathetic innervation and mammary gland ducts in more detail, we performed confocal imaging in six regions of the mgWAT fat pad from each of 6 RT and 6 cold-treated mice. Consistent with our scRNA-seq data, we saw a significant increase in EPCAM staining in the ducts of cold-exposed mgWAT (Fig 2C and 2D). We also identified contacts between TH+ fibres and EPCAM+ ducts (neuroductal points) with a trend to increased volume of nerve contacts (normalized for duct volume) in cold-treated mice (p = 0.09, Fig S2F). Interestingly, TH intensity, which has been reported to increase with sympathetic activation, was significantly higher at the neuroductal points in cold exposed mgWAT compared to controls (Fig 2C, 2E, 2F Supplementary Movie 2). Similarly, EPCAM intensity was also significantly elevated at neuroductal points in keeping with local induction of expression. However, there were no significant changes in duct or nerve volume or in TH intensity across the whole nerve volume (Extended Data Fig 2C–2H). Taken together, our data show a significant remodelling of mammary ducts and their contacts with sympathetic innervation upon cold exposure.

**Figure 2.**
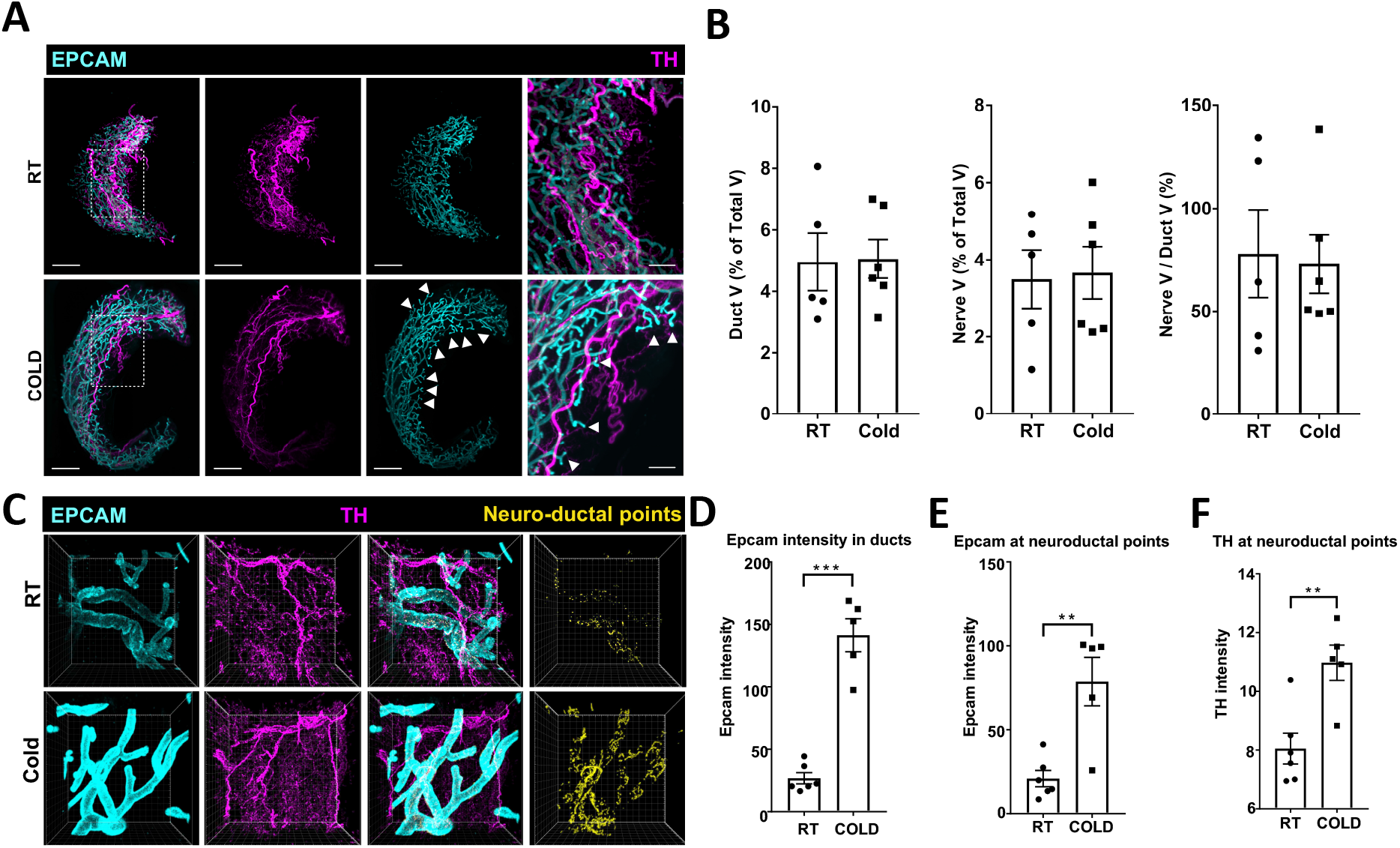
SNS fibers directly innervate mammary ductal epithelium. **(A)** Light sheet microscopy fluorescence (LSFM) images of mgWAT isolated from female mice exposed to RT or COLD for 24 hr and stained with TH antibody (SNS fibers) and EPCAM antibody (ductal cells). Representative mgWAT images from 5-6 mice per condition. White arrows show terminal ductal bifurcations under COLD condition **(B)** Quantification of LSFM images for ductal volume (Duct V) and nerve volume (Nerve V) as a percentage of total volume (Total V), and ratio of Nerve V and Duct V in RT or COLD mgWATs N=5-6 per condition. **(C)** Confocal images of mgWAT isolated from female mice exposed to RT or COLD (24 hr) and stained for EPCAM and TH antibodies. Merged stainings of EPCAM and TH represent neuroductal points. Representative image of 5-6 mice per condition. **(D-F)** Quantification of EPCAM intensity in ducts (D), EPCAM intensity at neuroductal points (E), and TH intensity at neuroductal points (F). N=5-6 per condition. **, p<0.01; ***, p<0.001.

To explore the biological relevance of cold-induced increase in i) luminal subtype population transcriptional state (Fig. 1) and ii) EPCAM intensity and SNS innervation of mammary ducts (Fig. 2), we first purified EPCAM+ and EPCAM-cells from mgWAT of RT and cold-exposed mice and then stained for EPCAM and CD49F for fluorescence-activated cell sorting (FACS) analysis to probe for luminal cell population changes under cold stress. As shown in Extended Data Fig. 3A, we noticed three distinct populations of cells that were EPCAM^lo^CD49F^lo^ (stromal), EPCAM^hi^CD49F^hi^ (luminal), and EPCAM^lo^CD49F^hi^ (basal) cells. Luminal cells were only enriched in EPCAM+ purified cells and showed a cold-dependent increase in cell population compared to RT conditions. We also noticed a marked reduction in the basal cell population upon cold treatment in EPCAM+ selected cells, consistent with the scRNA-seq data in Fig. 1C (myoepithelial cluster, RT: 2.15%, COLD: 0.65%). These data indicate a differential response to cold stress by luminal and basal cells in ducal epithelium (Extended Data Fig. 3A). To test whether luminal cells directly responded to SNS activation, we tested mammokine expression in isolated primary EPCAM+ mgWAT that were either treated or not treated with isoproterenol. Cold-induced mammokines showed increased expression upon isoproterenol (beta-adrenergic receptor agonist) treatment (Extended Data Fig. 3B). To determine whether adrenergic-activated luminal cells are involved in mgWAT adipose thermogenesis, we depleted EPCAM+ cells from mgWAT SVFs by positive selection using magnetic cell sorting (MACS) and differentiated SVFs *ex vivo* with and without ductal cells into beige adipocytes (Fig. 3A). Depletion of EPCAM+ cells from SVFs of RT mgWAT potentiated expression of thermogenic genes such as *Ucp1*, *Cox8b*, *Ppargc1a* and *Cidea* and this potentiation of thermogenic genes was markedly amplified in the cold-exposed condition (Fig. 3A and Extended Data Fig 3C). To further test the crosstalk of epithelial cells and adipocytes, we developed an *in vitro* co-culture system involving a controlled mixture of adipogenic 10T1/2 cells with nontransformed mouse mammary gland (NMuMG cells) (derived from “normal” mammary epithelium). Seeding density of as low as 2.5% NMuMG cells resulted in a significant reduction of *Ucp1* and beiging potential of 10T1/2 cells compared to the pure culture (Fig. 3B and Extended Data Fig. 3D–3F). The higher the fraction of NMuMG cells in the co-culture, the lower the relative expression of *Ucp1* and other thermogenic genes such as *Cox8B* and *Ppargc1a* measured by RT-qPCR (Fig. 3B). To test the *ex vivo* and *in vivo* role of ductal epithelial cells in adipose thermogenesis, we compared aged-matched female mgWAT with three complimentary ductal ablated models; i) estrogen receptor-alpha (ERa) knockout mice (*Esr1* KO), ii) male mice which lack or possess only rudimentary glandular ducts, iii) inguinal (ducts) or dorsolumbar (no ducts) portion of mgWAT from 5-week-old female mice. First, to test the role of ductal cells in adipose thermogenesis, we isolated SVFs from male iWATs and female mgWATs and differentiated them into beige adipocytes in the presence or absence of isoproterenol. We found that SVFs from male iWATs show markedly higher beiging and isoproterenol-mediated *Ucp1* expression compared to EPCAM+ SVFs from female mgWAT (Fig. 3C and Extended Data Fig. 3G). In agreement with the ex vivo data, female mgWATs showed significantly less expression of cold-induced thermogenic genes such as *Ucp1, Cox8b*, and *Ppargc1a* compared to male iWATs (Fig. 3D). Since mgWATs make up almost all of the subcutaneous fat mass in female mice, we reasoned that highly reduced adipocyte thermogenic gene expression could potentially influence whole body energy metabolism. To test this, we performed indirect calorimetry on age-matched male and female mice at RT and 24 hr cold exposure using a metabolic chamber. We found that female mice showed highly reduced energy expenditure (EE), oxygen consumption (VO2), and carbon dioxide production (VCO2) during cold exposure compared to male mice (Fig. 3E and Extended Data Fig. 3I). Female mice showed markedly higher respiratory exchange ratio (RER) than males under cold exposure, indicating a decrease in cold-induced fat oxidation and the possibility that mgWATs maintain adiposity under cold stress (Fig. 3E and Extended Data Fig. 3J). Generalized linear model (GLM)-based regression analyses showed a significant group and interaction effect in RER between males and females based on fat mass as a covariate (Extended Data Fig. 3J). We did not see significant differences in locomotor activities and food consumption between the sexes (Extended Data Fig. 3K). These RER data were further supported by our magnetic resonance imaging (MRI) body composition analysis which showed that male mice lose significant body weight and fat mass during cold stress whereas females show no differences before and after cold stress (Fig.3F and Extended Data Fig. 3H). Next, we compared the cold-induced thermogenic gene expression program between WT and Estrogen Receptor-alpha (*Esr1* KO) mice. *Esr1* KO mice are known to possess hypoplastic mammary ducts and remain rudimentary throughout the life span of a female mouse^36^. As shown in Fig. 3G, cold-exposed *Esr1* KO mice showed markedly increased expression of *Ucp1*, and other thermogenic genes compared to WT control. In the 5 weeks after birth, mammary ducts are concentrated in the inguinal portion closer to the nipple and are confined near the lymph node and virtually absent toward the dorsolumbar region of the mgWAT, providing distinct anatomical regions within the mgWAT to test the role of ductal epithelium in adipose thermogenesis^37^. 5-week-old female mice were exposed to cold and inguinal and dorsolumbar regions were dissected to assess thermogenic transcripts. *Epcam* transcripts were present only in the inguinal region, however, thermogenic genes were mostly similar between inguinal and dorsolumbar region except for *Ucp1* where we saw an increase in *Ucp1* levels in inguinal part (Fig. 3H). Chi et. al reported regional differences between inguinal and dorsolumbar region and there could be a regional control of *Ucp1* expression in 5-week-old mice independent of ductal cells^38^. However, consistent with our data, we noticed significantly less expression of genes involved in lipid mobilization such as *Pnpla2* and *Hsl*, in inguinal regions compared to dorsolumbar, indicating that the ductal epithelium is potentially inhibiting cold-induced lipid mobilization. All these data point, for the first time, toward a possible unique mechanism of SNS-activated mgWAT under cold stress to limit thermogenesis and preserve adiposity.

**Figure 3.**
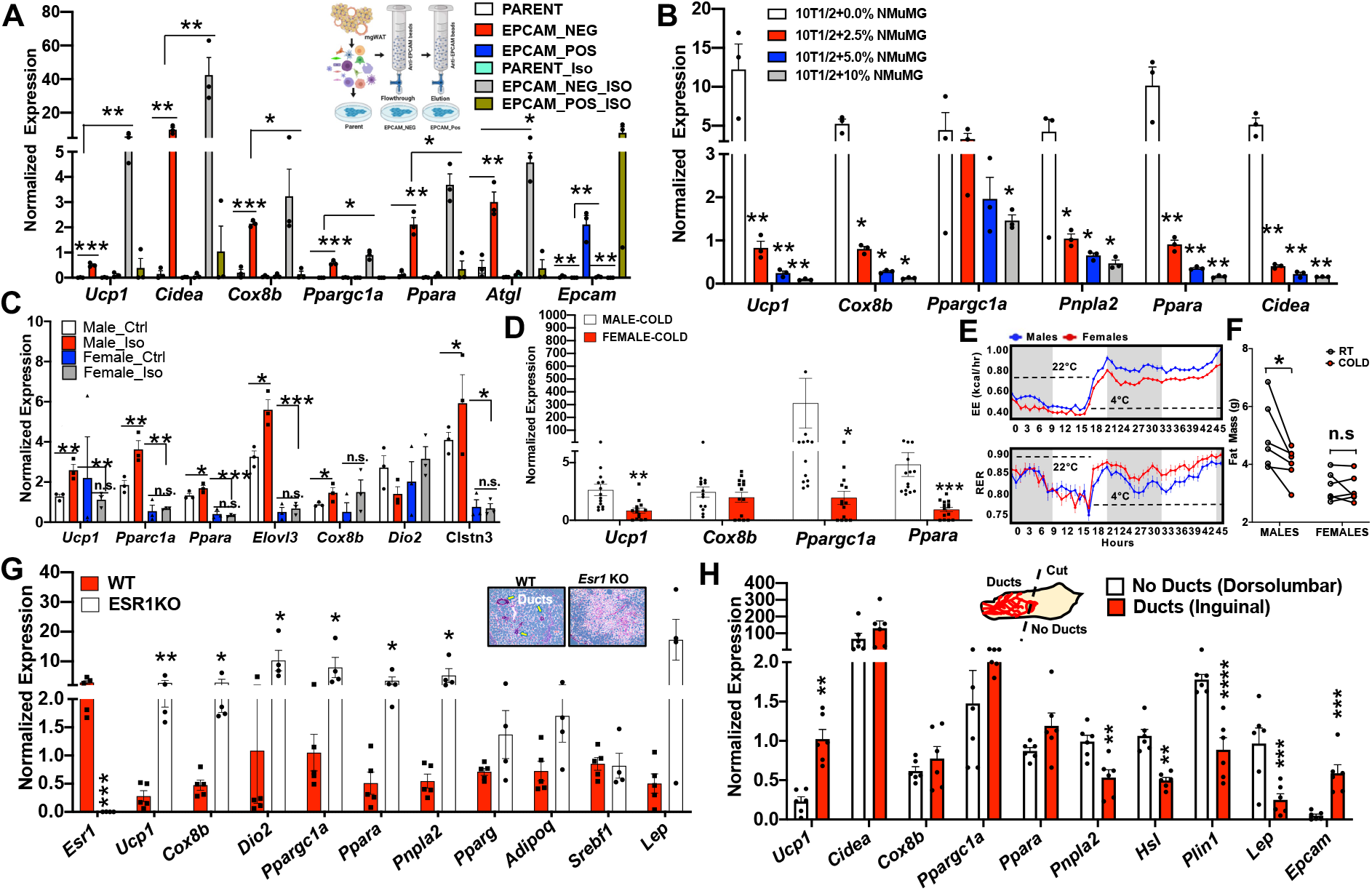
Mammary ductal cells directly inhibit adipocyte thermogenesis. **A)** Real-time qPCR of indicated genes from beige differentiation of primary mgWAT SVF (Parent), EPCAM-ve (EPCAM-NEG), and EPCAM+ (EPCAM-POS) cells treated with and without 10 μM isoproterenol (ISO) for 24 hr. Results are from three independent experiments. *, p<0.05; **, p<0.01; ***, p<0.001. Inset represent a cartoon depiction of selecting and plating EPCAM+ cells from SVFs derived from mgWATs. **B)** Real-time qPCR of indicated genes from beige differentiated and Iso treated 10T1/2 and 2.5-10% NMuMG mixture cells. Results are from three independent experiments. *, p<0.05; **, p<0.01. **C)** Real-time qPCR of indicated genes from beige differentiated SVFs isolated from male and female iWATs treated with and without Iso. Results are from three independent experiments. *, p<0.05, **, p<0.01; ***, p<0.001. **D)** Real-time qPCR of indicated genes from 24 h cold exposed male and female iWATs. N=13,13 *, p<0.05, **, p<0.01; ***, p<0.001. **E)** Energy expenditure (kcal/hr) and respiratory exchange ratio (RER) of male and female mice exposed to 22°C (21 hr) and 4°C (24 hr) analyzed in Sable Promethion metabolic chambers (12 hr light/dark cycle, 45 hr total duration, white bar represent light cycle and grey bar represent night cycle). N=6,6. **F)** Fat mass of mice from (E) at RT (22°C) and COLD (4°C). N=6,6. *, p<0.05; n.s, not significant. **G)** Real-time qPCR of indicated genes from 24 hr cold exposed WT and *Esr1* KO mgWATs. Inset picture shows histological section of mgWATs from WT and *Esr1* KO mice. Arrows are pointing towards ducts in WT. N=5,5 *, p<0.05, **, p<0.01. **H)** Real-time qPCR of indicated genes from dorsolumbar or inguinal parts of cold-exposed 5-week-old mice. Inset picture shows cartoon depiction of ducts (inguinal) and no ducts (dorsolumbar). Dotted line represents cut site to separate inguinal and dorsolumbar regions of mgWAT. N=6,6. *, p<0.05, **, p<0.01; ***, p<0.001

Our results show a unique possible SNS-mediated crosstalk between mammary ductal cells and adipocytes to control adipocyte thermogenesis. Our differentially expressed genes in mammary ductal cells under adrenergic stimulation showed upregulation of genes that encode factors such as Angptl4, Enho, Lrg1, and Lcn2, which are known to play inhibitory roles in adipocyte thermogenesis. These factors also showed high enrichment in mammary *Epcam+* cells by scRNA-seq and RNAscope in situ hybridization. Among these factors, *Lcn2* was previously shown to inversely correlate with *Ucp1* expression in female gonadal WAT (gWAT)^24^. Female gWATs, which lack mammary ducts, express exceedingly high levels of *Ucp1* compared to male gWATs^24^. Reciprocally, we find that cold-stressed female mgWATs express high levels of *Lcn2* and low levels of *Ucp1* compared to males (Fig. 3D and Extended Data Fig. 4A). The secretion of Lcn2 by luminal AV and HS-AV cells could potentially be a mechanism of luminal cells to block excess thermogenesis and preserve adiposity. Consistent with data in Fig.1 and Fig.3, both isoproterenol treatment and cold exposure led to an increase in *Lcn2* levels in *Epcam+* mammary ducts (Extended Data Fig. 4B). To test adrenergic-dependent expression of Lcn2 protein expression in luminal cells, we used organoids from a genetic mouse model that is based on doxycycline (Dox)-based tet-responsive diphtheria toxin A (DTA) system derived from interbreeding two transgenic strains: (1) mice expressing the tetracycline-on (“tet-on”) transcription factor rtTA under the control of the luminal epithelial cell-specific *Krt8* gene promoter (K8rtTA); (2) mice expressing tet-responsive DTA (TetO-Cre) that can be activated in the presence of Dox. This model enables luminal cell ablation in a temporally regulated manner. Mammary duct organoids isolated from *K8*rtTA-DTA mice^39^ show increases in Lcn2 protein expression upon isoproterenol treatment and these increases were diminished upon Dox treatment (Fig. 4A and Extended Fig. 4C–4H), showing a direct response of luminal cells to produce Lcn2. To test the physiological role of Lcn2 in regulating mgWAT thermogenesis, we mimicked cold induction of Lcn2 production in mgWAT by inducing Lcn2 exogenous expression specifically in mgWAT of *Lcn2* KO mice by injecting adipoAAV-Lcn2 or adipoAAV-GFP^24^ (See Methods). Our unbiased bulk RNA-seq data from the mgWATs of adipoAAV-Lcn2 injected mice show that *Lcn2* expression was not supraphysiological compared to controls (Fig. 4B). The volcano plot in Fig. 4B demonstrates that Lcn2 exogenous expression significantly decreased the expression of thermogenic genes such as *Ucp1, Cidea, Ppara* and increased expression of adipogenic genes including *Lep*, *Mmp12*^40,41^, *Zfp423*^42^, and Lbp^43^. Lcn2 overexpression also led to an increase in *Aldh1a1*, which was recently shown to inhibit adipose thermogenesis by downregulating UCP1 levels^44,45^. We validated our RNA-seq data by directed qPCR in both Lcn2 reconstituted mgWAT of *Lcn2* KO and WT mice (Fig. 4C) and treated beige differentiated SVFs isolated from *Lcn2* KO mgWATs with and without recombinant Lcn2 (Extended Data Fig. 4I). Finally, to test the role of Lcn2 under cold stress, female *Lcn2* KO and WT mice were exposed to cold (4°C) for 24 hr. Compared to controls, cold-exposed *Lcn2* KO mice mgWAT showed more beiging/browning and gene expression analysis showed a significant increase in thermogenic genes such as *Ucp1* indicating that Lcn2 is potentially one of the limiting factors involved in regulating the propensity of mgWAT to beige (Extended Data Fig. 4J). Overall, our scRNA-seq analysis and both our tissue-specific gain-of-function and loss-of-function experimental data show Lcn2 as a factor expressed in luminal cells that could function to inhibit thermogenesis and maintain adiposity in mgWAT during cold-stress.

**Figure 4.**
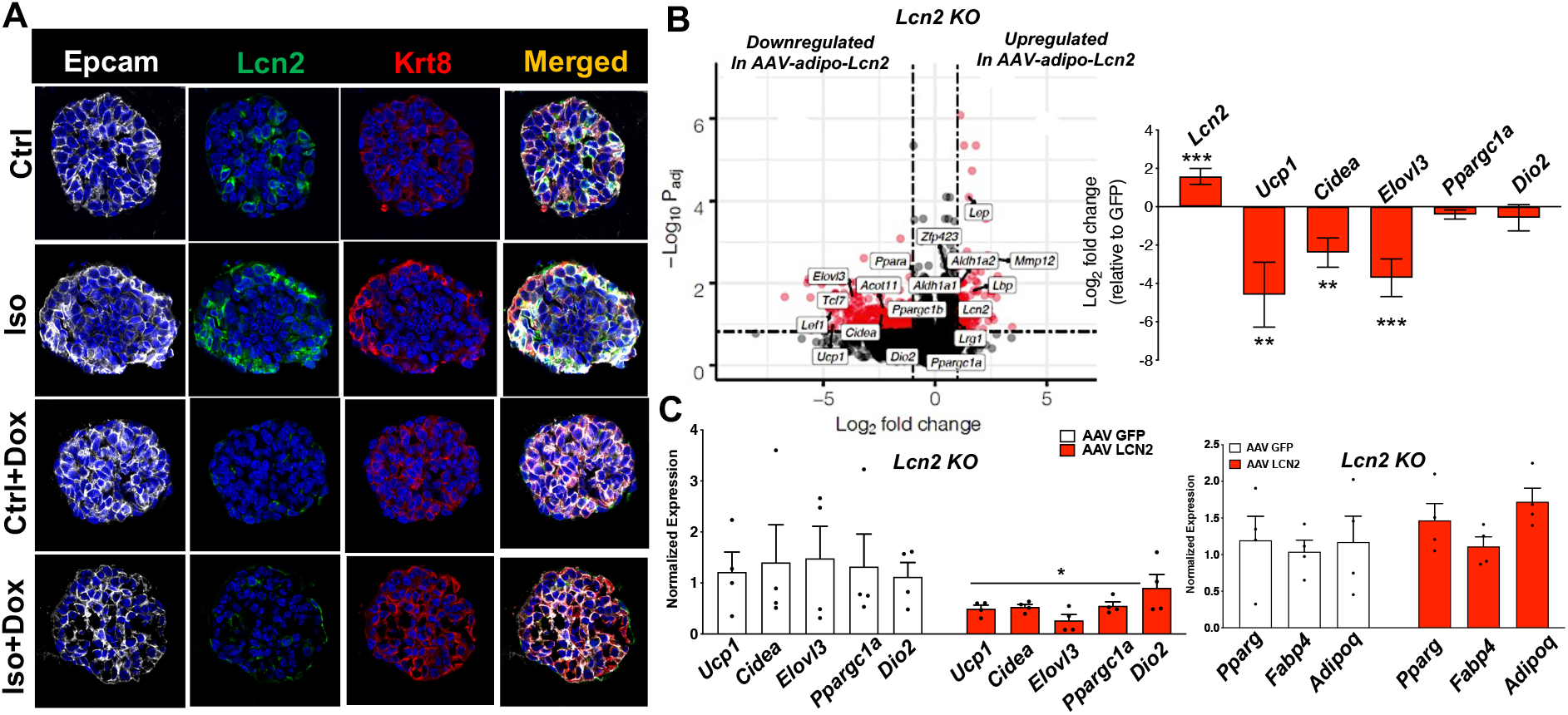
LCN2 preserves mgWAT adiposity. **A)** Confocal images of immunostaining of indicated antibodies in the organoids derived from K8rtTA-DTA mice (see Methods) treated with and without Iso and Doxycycline (Dox). Representative images from 3 organoid experiments. **B)** Volcano plot of DEGs from the mgWAT of *Lcn2* KO mice treated with adipose-specific AAV-LCN2 or AAV-GFP and represented as a fold change of LCN2/GFP ratio as a function of p-value. Genes labelled are either induced (+) or repressed (-) by Lcn2. **, p<0.01; ***, p<0.001. N=4,4. **C)** Real-time qPCR of indicated genes from the mgWATs of LCN2KO or WT mice treated with AAV-LCN2 or AAV-GFP. N=4,4. *, p<0.05

In summary, our studies have uncovered a direct inhibitory role of mammary duct luminal cells in adipocyte *Ucp1* expression and thermogenic gene program. SNS fibers directly innervate EPCAM+ luminal cells and adrenergic stimulation of luminal cells transduce expression of mammokines such LCN2 that regulates *Ucp1* expression in mgWAT adipocytes under cold exposure. Depletion of EPCAM+ luminal epithelial cells potentiate the capacity of ex vivo differentiated mgWAT preadipocytes to express UCP1, and female mice with ductal epithelium loss show higher cold-induced UCP1 expression compared to controls. Importantly, female mice demonstrate significantly less scWAT adipocyte UCP1 expression compared to male mice under cold exposure. Taken together, these findings provide a new insight into mammary gland biology, expand our understanding of the role of adipose microenvironment in adipocyte UCP1 expression, and reveal the potential of mammokines to regulate local and systemic energy homeostasis.

## METHODS

### Animal Studies

C57BL/6 WT male and female mice (#000664), *ESR1* KO (#004744), LCN2KO (#24630) were acquired from Jackson Laboratory and maintained in a pathogen-free barrier-protected environment (12:12 h light/dark cycle, 22°C-24°C) at the UCLA and Mount Sinai animal facilities. The Krt8rtTA-TetO-DTA mouse model was described previously^39^. For the time course cold exposure experiment, WT mice at 8-10 weeks of age were singly housed at 4°C room in a non-bedded cage without food and water for first 6 h; thereafter food, water, and one cotton square were added. For the 24 h harvest, 3 h before harvest, food, water, and cotton square were removed and then mice were harvested. At the end of the experiment, mgWATs were resected for analysis. For overexpression studies, recombinant adeno-associated virus serotype 8 (AAV8) expressing LCN2 or GFP was generated and injected as described previously^24^. Indirect calorimetry was performed using Promethion Systems (Sable Systems, Las Vegas, NV). Animals were placed individually in chambers at ambient temperature (22.0 °C) for 21 hr followed by 24 hr cold (4.0 °C) with 12 hr light/dark cycles. Animals had free access to food and water. Respiratory measurements were made in 5 min intervals after initial 7-9 hr acclimation period. Energy expenditure was calculated from VO2 and RER using the Lusk equation, EE in Kcal/hr = (3.815 + 1.232 X RER) X VO2 in ml/min. Indirect calorimetry data were analyzed by CALR web-based software^46^. Body composition (fat mass) was determined using EchoMRI Body Composition Analyzer. Animal experiments were conducted in accordance with the Mount Sinai and UCLA Institutional Animal Care and Research Advisory Committee.

### RNA-Seq

RNA isolation, library preparation, and analysis were conducted as previously described^24^. Flash-frozen mgWAT samples were homogenized in QIAzol (Qiagen, Germantown, MD), and following chloroform phase separation, RNA was isolated according to the manufacturer’s protocol using miRNeasy columns (Qiagen, Germantown, MD). Libraries were prepared from extracted mgWAT fat RNA (Agilent 2200 Tapestation eRIN >8.2) using KAPA Stranded mRNA-Seq Kit (cat #KK8421, KAPA Biosystems, Wilmington, MA), per the manufacturers’ instructions. The pooled libraries were sequenced using an Illumina HiSeq4000 instrument with SE50bp reads (Illumina, San Diego, CA). Reads were aligned to the mouse genome mm10 using STAR^47^ or HISAT2^48^ aligner and quantified using the Bioconductor R packages as described in the RNA-Seq workflow^49^. *P* values were adjusted using the Benjamini-Hochberg procedure of multiple hypothesis testing^49^.

### Single cell isolation from SVF

Single cell SVF populations from adipose tissue were isolated as described previously^3,31^. The fourth inguinal white adipose tissue (iWAT) depot mgWAT from mice exposed to cold stress (4°C) or room temperature for 24 hr was dissected and placed on a sterile 6-well tissue culture plate with ice-cold 1X DPBS. Excess liquid was removed from fat pads by blotting. Each tissue was cut and minced with scissors and then placed in 15 ml conical tubes containing digestion buffer (2 ml DPBS and Collagenase II at 3 mg/ml; Worthington Biochemical, Lakewood, NJ, USA) for 40 min of incubation at 37°C with gentle shaking at 100 rpm. Following tissue digestion, enzyme activity was stopped with 8 ml of resuspension media (DMEM/F12 with glutamax supplemented with 15%FBS and 1% pen/strep; Thermo Scientific, CA). The digestion mixture was passed through 100 μm cell strainer and centrifuged at 150 x g for 8 min at room temperature. To remove red blood cells, the pellet was resuspended and incubated in RBC lysis buffer (Thermo Scientific, CA) for 3 min at room temperature, followed by centrifugation at 150 x g for 8 min. The pellet was then resuspended in resuspension media and again spun down at 150 x g for 8 min. The cell pellet was resuspended in 1 ml of 0.01% BSA (in DPBS) and passed through a 40 μm cell strainer (Fisher Scientific, Hampton, NH, USA) to discard debris. Cell number was counted for 10X Genomics single cell application.

### SVF single cell barcoding and library preparation

To yield an expected recovery of 4000-7000 single cells, an estimated 10,000 single cells per channel were loaded onto Single Cell 3’ Chip (10X Genomics, CA). The Single Cell 3’ Chip was placed on a 10X Genomics instrument to generate single cell gel beads in emulsion (GEMs). Chromium Single Cell 3’ v3 Library and Cell Bead Kits were used according to the manufacturer’s instructions to prepare single cell RNA-Seq libraries.

### Illumina high-throughput sequencing libraries

Qubit Fluorometric Quantitation (ThermoFisher, Canoga Park, CA, USA) was used to quantify the 10X Genomics library molar concentration and a TapeStation (Aligent, Santa Clara, CA, USA) was used to estimated library fragment length. Libraires were pooled and sequenced on an Illumina HiSeq 4000 (Illumina, San Diego, CA, USA) with PE100 reads and an 8 bp index read for multiplexing. Read 1 contained the cell barcode and UMI and read 2 contained the single cell transcripts.

### Single cell data pre-processing and quality control

To obtain digital gene expression matrices (DGEs) in sparse matrix representation, paired end reads from the Illumina HiSeq 4000 were processed and mapped to the mm10 mouse genome using 10X Genomics’ Cell Ranger v3.0.2 software suite. Briefly, .bcl files from the UCLA Broad Stem Cell Research Center sequencing core were demultiplexed and converted to fastq format using the ‘mkfastq’ function from Cell Ranger. Next, the Cell Ranger ‘counts’ function mapped reads from fastq files to the mm10 reference genome and tagged mapped reads as either exonic, intronic, or intergenic. Only reads which aligned to exonic regions were used in the resulting DGEs. After combining all four sample DGEs into a single study DGE, we filtered out cells with (1) UMI counts < 700 or > 30,000, (2) gene counts < 200 or > 8,000, and (3) mitochondrial gene ratio > 10%. This filtering resulted in a dataset consisting of 42,052 genes across 12,222 cells, with approximately 2,300 – 4,650 cells from each sample. A median of 2,411 genes and 7,252 transcripts were detected per cell.

### Identification of cell clusters

To achieve high resolution cell type identification and increased confidence in our cell type clustering we brought in external publicly available single cell data from SVF and mammary tissues. Specifically, we included single cell data from 9 datasets comprising 91,577 single cells from the mammary gland and multiple adipose depots, across 4 different single cell platforms (**Table 1**). These external datasets and the SVF data from this study were all independently normalized using sctransform^50^ and integrated using Seurat^51,52^ v3.1.5. The single cell expression profiles were projected into two dimensions using UMAP^53^ or tSNE^54^ and the Louvain^55^ method for community detection was used to assign clusters. This integrated data was only used to identify and define the cell types. All plots which are not explicitly designated as integrated with at least one external dataset and all downstream analyses (e.g. differential expression analyses) were conducted on non-integrated data to retain the biological effect of the cold treatment. Visualization of the non-integrated data was conducted on a subsampled dataset where all samples had the same number of cells to give an equal weight to each sample, however, all downstream analyses (e.g. differential expression analyses) leveraged the full dataset.

### Cell type-specific gene expression signatures

Cell type-specific gene expression signatures were generated by identifying genes with expression levels two-fold greater (adjusted p-values < 0.05) than all other cell types. To ensure consistency across samples, Seurat’s FindConservedMarkers function (Wilcoxon rank sum test with a meta p-value) was applied across each sample.

### Resolving cell identities of the cell clusters

To identify the cell type identity of each cluster, we used a curated set of canonical marker genes derived from the literature (**Supplementary Table 1**) to find distinct expression patterns in the cell clusters. Clusters which uniquely expressed known marker genes were used as evidence to identify that cell type. Cell subtypes which did not express previously established markers were identified by general cell type markers and novel markers obtained with Seurat’s FindConservedMarkers function were used to define the cell subtype.

### Differential gene expression analysis

Within each identified cell type, cold treated and room temperature single cells were compared for differential gene expression using Seurat’s FindMarkers function (Wilcoxon rank sum test) in a manner similar to Li et al.^19^. Differentially expressed genes were identified using two criteria: (i) an expression difference of >= 1.5-fold and adjusted p-value < 0.05 in a grouped analysis between room temperature mice (n = 2) and cold treated mice (n = 2); (ii) an expression difference of >= 1.25 fold and consistent fold change direction in all 4 possible pairwise combinations of cold-treated vs room temperature mice.

### Pathway enrichment analysis

Pathway enrichment analysis was conducted on the differentially expressed genes from each cell type using gene sets from KEGG^62^, Reactome^63^, BIOCARTA^64^, GO Molecular Functions^65^, and GO Biological Processes^65^. Prior to enrichment, mouse gene names were converted to human orthologues. Enrichment of pathways was assessed with a Fisher’s exact test, followed by multiple testing correction with the Benjamini-Hochberg method. Gene set enrichments with FDR < 0.05 were considered statistically significant.

### Real time qPCR

Total RNA was isolated using TRIzol reagent (Invitrogen) and reverse transcribed with the iScript cDNA synthesis kit (Biorad). cDNA was quantified by real-time PCR using SYBR Green Master Mix (Diagenode) on a QuantStudio 6 instrument (Themo Scientific, CA). Gene expression levels were determined by using a standard curve. Each gene was normalized to the housekeeping gene 36B4 and was analyzed in duplicate. Primers used for real-time PCR are previously described^3,31^ and presented in Table 2.

### RNAScope Fluorescence in situ hybridization (FISH)

mgWAT from RT or cold exposed mice (Jackson Laboratory, #000664) was fixed in 10% formalin overnight, embedded with paraffin, and sectioned into unstained, 5 μm-thick sections. Sections were baked at 60°C for 1 hour, deparaffinized, and baked again at 60°C for another hour prior to pre-treatment. The standard pre-treatment protocol was followed for all sectioned tissues. *In situ* hybridization was performed according to manufacturer’s instructions using the RNAscope Multiplex Fluorescent Reagent Kit v2 (#323136, Advanced Cell Diagnostics [ACD], Newark, CA). Opal fluorophore reagent packs (Akoya Biosciences, Menlo Park, CA) for Opal 520 (FP1487A), Opal 570 (FP1488A), Opal 620 (FP1495A), and Opal 690 (FP1497A) were used at a 1:1000 dilution in TSA buffer (#322809, ACD). RNAscope probes from ACD were used for the following targets: EPCAM (#418151), ENHO (#873251), LRG1 (#423381), LCN2 (#313971), HP (#532711), WNT4 (#401101), NRG4 (#493731), KRT8 (#424528), and MFGE8 (#408778). Slides were mounted with ProLong Diamond Antifade Mountant with DAPI (P36962, Life Technologies, Carlsbad, CA). Fluorescent signals were captured with the 40x objective lens on a laser scanning confocal microscope LSM880, (Zeiss, White Plains, NY).

### Fluorescent-activated cell sorting (FACS)

Mammary gland white adipose tissue (mgWAT) from RT or cold exposed mice (Jackson Laboratory, #000664) was dissected, cut, minced, and digested with collagenase D (5 mg/mL, #11088882001, Roche, Germany) and dispase (2 mg/mL, #17105041, Gibco, Grand Island, NY) over 40 min at 37°C with gentle shaking at 100 RPM. Enzymatic digestion was stopped with DMEM/15% FBS and the cell suspension was filtered through a 100 μm nylon mesh cell strainer, and centrifuged for 10 minutes at 700 x g. SVF pellet was resuspended in 1 mL Red Blood Cell lysis buffer (#41027700, Roche, Germany) and incubated for 5 minutes at room temperature. Cell suspension was diluted in 4 mL DPBS and filtered through a 40 μm nylon mesh cell strainer and centrifuged for 10 minutes at 700 x g. Single cell suspension was blocked for 10 minutes on ice in 500 μL DPBS/5% BSA (blocking buffer), centrifuged for 10 min at 700 x g, resuspended in 200 μL of DBPS/0.5% BSA (FACS buffer) solution containing the desired antibody mix, and incubated for 1 hour at 4°C in the dark with gentle rotation. Antibody-stained samples were washed with 800 μL FACS buffer, centrifuged 10 minutes at 700 x g, and resuspended in FACS buffer containing DAPI (at 1 ug/mL). Flow cytometry analysis was performed on a BD FACS Canto II (BD Biosciences, San Jose, CA) and results analyzed on FCS Express software (DeNovo Software, Pasadena, CA). Fluorescently-tagged anti-mouse antibodies (BioLegend, San Diego, CA) were used to label cell surface markers for flow cytometry analysis: EPCAM-FITC (clone G8.8, #118207), Sca-1-APC (Ly6, clone E13-161.7, #122512), CD49f-APC (clone GoH3, #313616). For flow cytometry analysis, negative selection of CD45-expressing cells using CD45 microbeads (#130052301) was performed immediately prior to the EPCAM positive selection protocol described above.

### Isolation, selection, and *ex vivo* treatment of EPCAM-positive cells

MACS microbeads (Miltenyi Biotec, Auburn, CA) were used for immuno-magnetic labeling positive selection of EPCAM-expressing cells (anti-CD326, #130105958). Before magnetic labeling, a single-cell suspension from the stromal vascular fraction of female mouse iWAT was prepared in MACS buffer, i.e. PBS, pH 7.2, 0.5% bovine serum albumin (#A7030, SIGMA, St. Louis, MO) and 2 mM EDTA, filtered through a MACS pre-separation 30 μm nylon mesh (#130041407) to remove cell clumps. Then, for magnetic labeling of EPCAM-expressing cells, 10 μL of EPCAM microbeads were added per 1×10^7^ total cells in 100 uL buffer, incubated for 15 minutes with rotation at 4°C, washed with 1 mL buffer, centrifuged at 700 x *g* for 5 minutes, resuspended in 500 μL buffer, and added to a pre-equilibrated MACS LS column (#130042401) in the magnetic field of a MACS separator (#130042302). Unlabeled EPCAM-negative cells were collected in the flow-through with three subsequent washes. The column was removed from the magnetic field, 5 mL of MACS buffer were added to the column, and the magnetically-labeled EPCAM-positive cells retained in the column were collected by flushing the cells down the column with a plunger. Finally, EPCAM-negative and EPCAM-positive cell populations were centrifuged at 700 x *g* for 5 minutes, resuspended in DMEM/F12 with glutamax supplemented with 15% FBS and 1% pen/strep (Thermo Scientific, CA) and plated on Collagen I-coated 12-well tissue culture plates (#354500, Corning, Kennebunk, ME). Media was replaced every other day during 6 days, followed by cell lysis with Tryzol for phenol/chloroform RNA extraction, and RT-qPCR analysis.

### Adipocyte differentiation and treatments

10T1/2 or SVF from the 4^th^ inguinal (iWAT) mgWAT was isolated from 8 week old female *Lcn2-* null mice, respectively. 10T1/2 cells were maintained as previously described^3^. The pre-iWAT cells were maintained in Dulbecco’s Modified Eagle Medium: Nutrient Mixture F-12 (DMEM/F12) supplemented with 1% glutamax, 10% fetal calf serum and 100 U/ml of both penicillin and streptomycin (basal media). Two days after plating (day 0), when the cells reached nearly 100% confluency, the cells were treated with an induction media containing basal media supplemented with 4 μg/mL insulin, 0.5 mM IBMX, 1 μM dexamethasone, and 1 μM rosiglitazone. After 48 h, the cells were treated with a maintenance media containing the basal media supplemented with 4 μg/mL insulin, and 1 μM rosiglitazone, with a media change every 2 days until day 10. For qPCR, differentiated iWAT cells were treated with 1 μg/ml recombinant LCN2 (Sino Biological Inc.) or differentiated 10T1/2 cells were treated with LRG1 (R&D Systems) for 24 h and then treated with isoproterenol (Sigma) for 6 h after which RNA was collected.

### iDISCO and Adipoclear tissue labelling and clearing

#### Sample collection

Immediately after cold exposure mice were anaesthetized with isoflurane (3%) and perfused with heparinized saline followed by 4% paraformaldefyde (PFA) (Electron Microscopy Sciences, Hatfield, PA, USA). Fat pads were carefully dissected and postfixed overnight in 4% PFA at 4°C. On the following day the tissue was washed 3 times in PBS before proceeding with the optical clearing protocol.

#### Optical clearing

Whole-mount staining and clearing was performed using the Adipo-Clear protocol as previously described^1^. Dissected fat pads were dehydrated at room temperature (RT) with a methanol/B1n buffer (0.3 M glycine, 0.1% Triton X-100 in H_2_O, pH 7) gradient (20%, 40%, 60%, 80%, 100%), incubated 3 times (1h, o/n, 2h) in 100% dichloromethane (DCM) (Sigma-Aldrich, St. Louis, MO, USA) to remove hydrophobic lipids, washed twice in 100% methanol, and bleached in 5% H_2_O_2_ overnight at 4°C to reduce tissue autofluorescence. Fat pads were then rehydrated with a methanol/B1n buffer gradient (80%, 60%, 40%, 20%) and then washed twice in 100% B1n buffer (1h, o/n). Primary antibodies (EPCAM: 1:250; TH: 1:500) were diluted in modified PTxwH (PBS with 0.5% Trixon X-100, 0.1% Tween-20, 2 μg/ml heparin, as previously described^66^ and applied for 6 days at 37°C. Following 5 washes with modified PTxwH over 1 day with the last wash performed overnight, secondary antibodies were diluted in modified PTxwH (1:500) and samples incubated for 6 days at 37°C. Samples were washed 5 times over 1 day in modified PTxwH at 37°C, 5 times over 1 day in PBS at RT, embedded in 1% agarose, dehydrated with a methanol gradient in H_2_O (12%, 50%, 75%, 100%), washed 3 times for 1 hr in 100% methanol followed by 3 times for 1 hr in DCM, before being transferred to dibenzylether (DBE) (Sigma-Aldrich) to clear.

#### Imaging

Z-stacked optical sections of whole fat pads were acquired with an Ultramicroscope II (LaVision BioTec, Bielefeld, Germany) at a 1.3x magnification with a 4 μm step size and dynamic focus with a maximum projection filter. Samples were then imaged in glass-bottom μ-dishes (81158, Ibidi, Gräfelfing, Germany) using an inverted Zeiss LSM 710 confocal microscope with a 10x (NA: 0.3) objective and a step size of 5 μm.

#### Image analysis

Imaris versions 9.6.0-9.7.2 (Bitplane AG, Zürich, Switzerland) were used to create digital surfaces covering ducts, TH+ innervation and total sample volume (1.3x light sheet images and 10x confocal images) to automatically determine volumes and intensity data. Volume reconstructions were performed using the surface function with local contrast background subtraction. For detection of EPCAM+ ducts in 1.3x light sheet images, a smoothing factor of 5 μm was used and the threshold factor was set to correspond to the largest duct diameter in each sample. For detection of TH+ nerves in 1.3x light sheet images, a smoothing factor of 3 μm and a threshold factor of 80 μm were used. For detection of EPCAM+ ducts in 10x confocal images, a smoothing factor of 3.35 μm was used and a threshold factor corresponding to the diameter of the thickest duct wall in each sample was used. For detection of TH+ nerves in 10x confocal images, a smoothing factor of 2 μm and a threshold factor of 5 μm were used. In 10x confocal images, nerve/duct interactions were defined by masking the TH channel using the TH+ nerve surface to remove any background, and then masking it again using the EPCAM+ duct surface. This process revealed TH+ innervation overlapping with EPCAM+ staining. A new surface was created to cover this overlapping TH+ innervation using a smoothing factor of 2 μm and a threshold factor of 5 μm.

#### Statistical analyses

Data are shown as mean±S.E.M. Distribution was assessed by Shapiro-Wilk test. Significance was determined by a two-tailed unpaired *t* test (parametric distribution) or by a Mann-Whitney test (non-parametric distribution). Significance was set at an alpha level of 0.05.

#### Mammary gland organoids culture

For organoid culture, we used a previously published protocol^67^. In brief, fat pads of 8–9-week-old K8rtTA/TetO-DTA^39^ virgin female mice were dissected and the lymph nodes removed. Tissues were briefly washed in 70% ethanol and manually chopped into 1 mm^3^ pieces. The finely minced tissue was transferred to a digestion mix consisting of serum-free Leibovitz’s L15 medium (Gibco) containing 3 mg ml^-1^ collagenase A (Sigma) and 1.5 mg ml^-1^ trypsin (Sigma). This was incubated for 1 hr at 37 °C to liberate epithelial tissue fragments (‘organoids’). Isolated organoids were mixed with 50 μl of phenol-red-free Matrigel (BD Biosciences) and seeded in 24-well plates. The basal culture medium contained phenol-red-free DMEM/F-12 with penicillin/streptomycin, 10 mM HEPES (Invitrogen), Glutamax (Invitrogen), N2 (Invitrogen) and B27 (Invitrogen). The basal medium was supplemented with Nrg1 (100 ng ml^-1^, R&D), Noggin (100 ng ml^-1^, Peprotech) and R-spondin 1 (100 ng ml^-1^, R&D). Then, 500 μl supplemented basal culture medium was added per well and organoids were maintained in a 37 °C humidified atmosphere under 5% CO_2_. After one week in culture, mammary organoids were released from the Matrigel by breaking the matrix with a P1000 pipette on ice. After 2–3 passages of washing and centrifugation at 1,500 rpm (140g) for 5 min at 4 °C, mammary cells were resuspended in Matrigel, seeded in 24-well plates and exposed to the previously described culture conditions. Organoids were treated either with 10 μg ml^-1^ of DOX to promote luminal cell ablation or with 10μM of isoproterenol (Sigma) for 6 hr. After 6 hr treatment, organoids were collected from Matrigel as mentioned before to perform further analysis.

#### RNA extraction and quantitative real-time PCR in organoid samples

To perform RNA extraction, isolated organoids were collected into kit lysis buffer. RNA was extracted with Qiagen RNeasy Micro Kit. After nanodrop RNA quantification and analysis of RNA integrity, purified RNA was used to synthesize the first-strand cDNA in a 30 μl final volume, using Superscript II (Invitrogen) and random hexamers (Roche). Genomic contamination was detected by performing the same procedure without reverse transcriptase. Quantitative PCR analyses were performed with 1 ng of cDNA as template, using FastStart Essential DNA green master (Roche) and a Light Cycler 96 (Roche) for real-time PCR system. Relative quantitative RNA was normalized using the housekeeping gene Gapdh. Analysis of the results was performed using Light Cycler 96 software (Roche) and relative quantification was performed using the ddCt method using Gapdh as a reference.

### Immunofluorescence in organoid samples

For immunofluorescence, collected organoids were pre-fixed in 4% PFA for 30 min at RT. Pre-fixed organoids were washed in 2%FBS-PBS, embedded in OCT and kept at −80°C. Sections of 4 μm were cut using a HM560 Microm cryostat (Mikron Instruments). Tissue sections were incubated in blocking buffer (BSA 1%, HS 5%, Triton-X 0.2% in PBS) for 1 hr at RT. The different primary antibodies were incubated overnight at 4 °C. Sections were then rinsed in PBS and incubated with the corresponding secondary antibodies diluted at 1:400 in blocking buffer for 1 hr at RT. The following primary antibodies were used: rat anti-K8 (1:1,000, Troma-I, Developmental Studies Hybridoma Bank, University of Iowa), rabbit anti-EPCAM (1:1,000, ab71916, Abcam), goat anti-Lcn2 (1:50, AF1857, R&D). The following secondary antibodies, diluted 1:400, were used: anti-goat (A11055) conjugated to Alexa Fluor 488 (Invitrogen), anti-rat (712-295-155) rhodamine Red-X and anti-rabbit (711-605-152) Cy5 (Jackson ImmunoResearch). Nuclei were stained with Hoechst solution (1:2,000) and slides were mounted in DAKO mounting medium supplemented with 2.5% DABCO (Sigma).

## Acknowledgements

A.A is supported by senior postdoctoral fellowship from the Charles H. Revson Foundation (grant no. 18-25), a fellowship from Sweden-America Foundation (Ernst O Eks fond), and a postdoctoral scholarship from the Swedish Society for Medical Research (SSMF). S.A.S is supported by American Diabetes Association Pathway to Stop Diabetes Grant ADA #1-17-ACE-31 NIH (R01NS097184, OT2OD024912, and R01DK124461) Department of Defense (W81XWH-20-1-0345, W81XWH-20-1-0156). A.J.L is supported by NIH U01 AG070959 and U54 DK120342. P.R is supported by R00DK114571, NIDDK-supported Einstein-Sinai Diabetes Research Center (DRC) Pilot & Feasibility Award, and Diabetes Research and Education Foundation (DREF) Grant # 501 (PR). The funders had no role in study design, data collection and interpretation, or the decision to submit the work for publication.

## Contributions

D.A. performed all the scRNA-seq data analysis under the supervision of X.Y. L.C.S. performed most of the biological experiments under the supervision of P.R. A.A. performed iDISCO and data analysis under the supervision of S.A.S. K.C.K performed LCN2 related animal experiments under the supervision of A.J.L. A.C.S performed organoid experiments under the supervision of C.B. S.P. performed RNAscope and EPCAM cells isolation experiments under supervision of L.C.S and P.R. S.S. performed indirect calorimetry and body composition studies under the supervision of P.R. I.S.A, G.D., and I.C., prepared single cell suspensions of mgWAT SVFs under the supervision of P.R. and X.Y. P.R. conceived the project and wrote the manuscript with help from A.J.B., C.B., S.A.S, A.J.L., and X.Y.

---

**Extended Data Figure 1.**
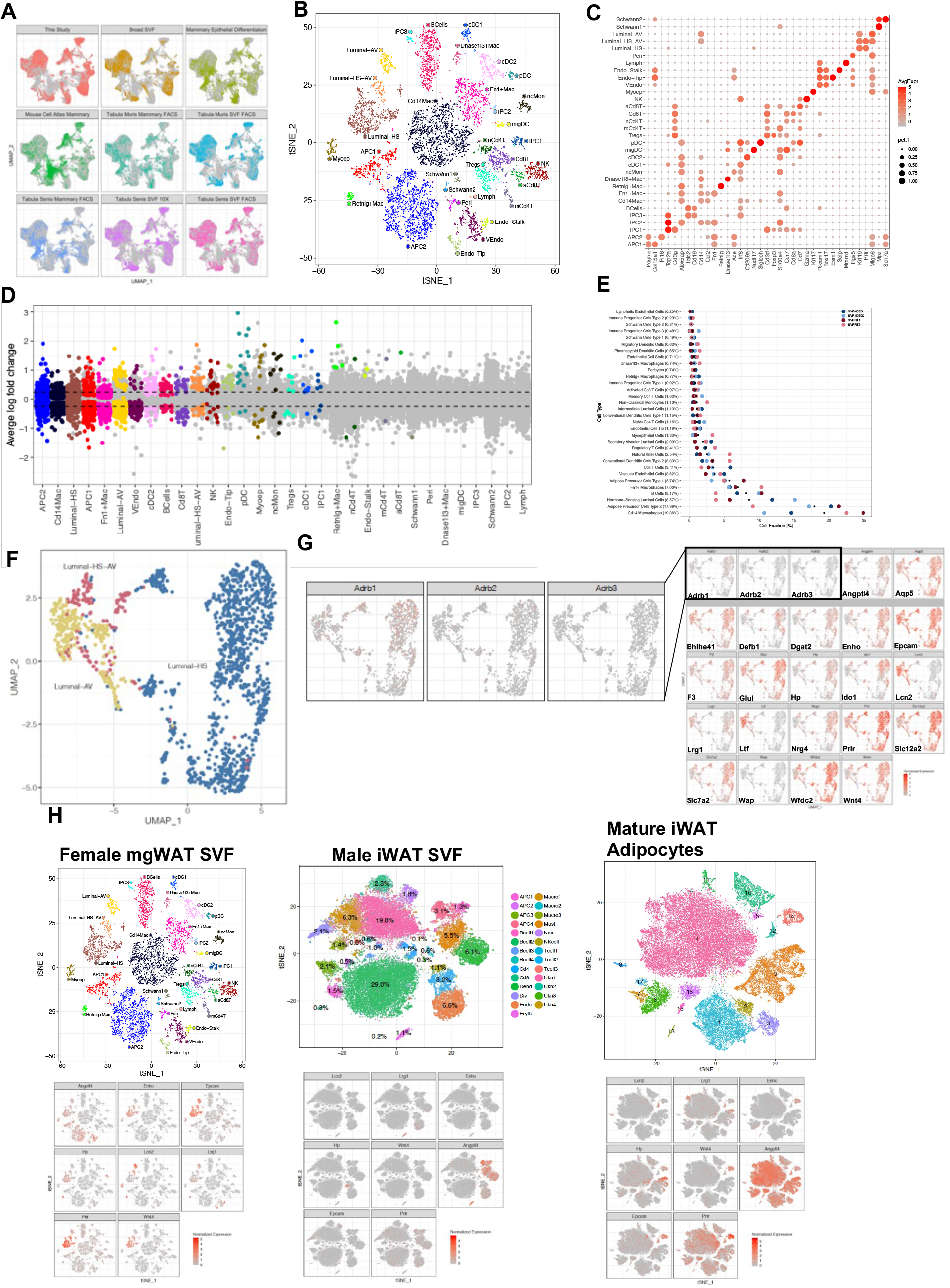
Cold-associated increase in cell percentages of luminal epithelium subtypes. **A)** UMAP plots of integrated single cell data from this study and 8 external datasets (see Methods and Table 1). Each point represents a single cell and are colored by dataset. **B)** t-SNE plot of single cells from mammary gland and surrounding SVF from this study colored by cell type. Cell types were identified based on expression of canonical marker genes. **C)** Expression of known canonical markers for cell types in the SVF and mammary gland. Color corresponds to average expression level and size corresponds to percentage of cells which express the gene within the cluster. **D)** Differentially expressed genes between COLD (24 hr) treated mice and RT animals across all cell types. Significant DEGs (adjusted p-value < 0.05) are highlighted with the average log fold change between 4 degree and RT indicated on the y-axis. Cell types are ordered on the x-axis based on the number of significant DEGs. **E)** Relative fractions of cell types within each sample. Black dots indicate average relative fractions across all samples. **F)** Aggregated UMAP plot of subclustering of luminal single cells from RT and cold-exposed mice. **G)** UMAP plots of normalized gene expression levels for genes of interest in luminal cells. **H)** tSNE plots of cell type clusters and normalized gene expression levels for genes of interest across multiple datasets including: female mgWAT SVFs, male iWAT SVFs and mature mgWAT adipocytes.

**Extended Data Figure 2.**
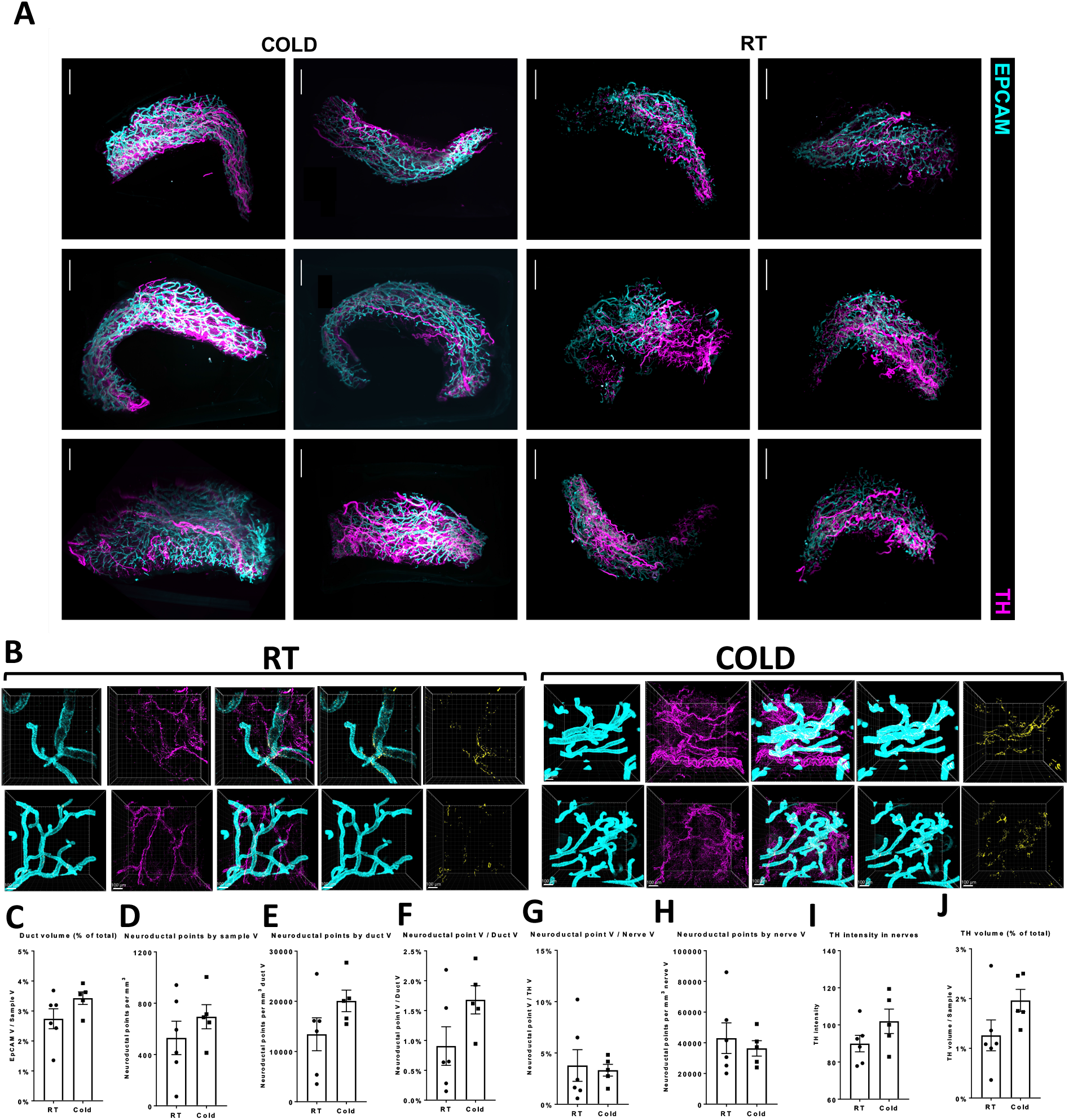
SNS fibers directly innervate ductal epithelial cells. **A)** Light sheet microscopy fluorescence (LSFM) images of mgWAT isolated from female mice exposed to RT or COLD for 24 hr and stained with TH antibody (SNS fibers) and EPCAM antibody (ductal cells). N=6,6. **B)** Confocal images of mgWAT isolated from female mice exposed to RT or COLD for 24hr and stained for EPCAM and TH antibodies. Merged staining of EPCAM and TH represent neuroductal points. Representative images of 5-6 mice per condition. **C-J)** Quantification of indicated parameters of images from (B). N=5-6 mice per condition.

**Extended Data Figure 3.**
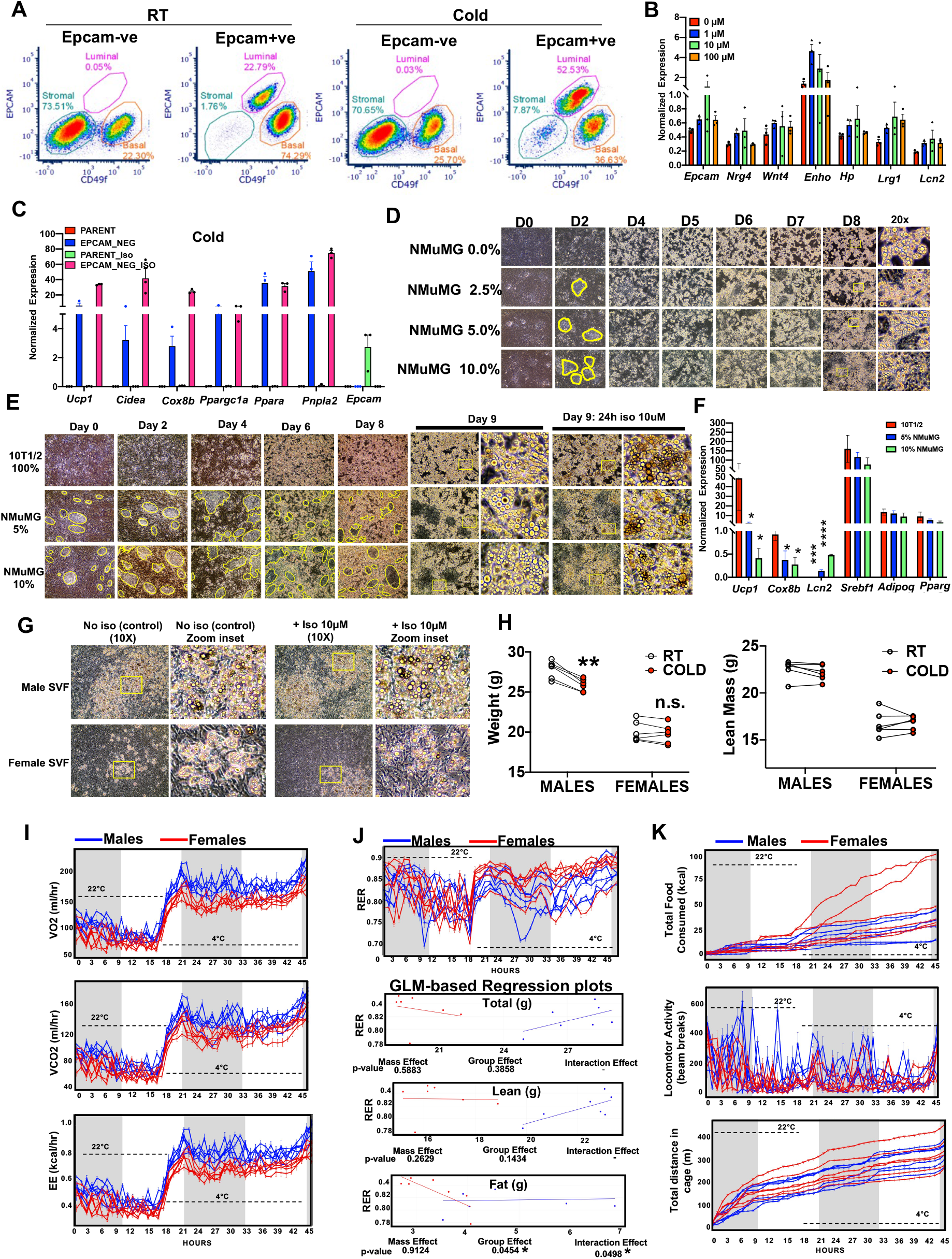
Mammary gland epithelium inhibits cold-induced adipocyte thermogensis. **A)** Representative FACS plot of CD49f and EPCAM expression in EPCAM bead selected EPCAM+ or EPCAM-ve epithelial cells from RT or cold mice. 2 mice per condition and 4 mammary fat pads per mouse. Representative data from 4 independent experiments. **B)** Real-time qPCR of indicated genes in beige differentiated SVFs from mgWAT treated with indicated concentration of Iso for 5 hr. **C)** Real-time qPCR of indicated genes from beige differentiated primary mgWAT SVF (Parent) and EPCAM-ve (EPCAM-NEG) cells isolated from cold exposed mice treated with and without 10 μM isoproterenol (ISO) for 24 hr ex vivo. Results are from three independent experiments. **D)** Images showing cell morphology of D0-D8 beige differentiated 10T1/2 and NMuMG (0, 5, and 10%) mixture cells. Yellow enclosures show NMuMG epithelial cells surrounded by 10T1/2 cells. **E)** Images showing cell morphology of D0-D9 beige differentiated 10T1/2 and NMuMG (0-10%) mixture cells. Yellow enclosures show NMuMG epithelial cells surrounded by 10T1/2 cells. **F)** Real-time qPCR of indicated genes from cells in (E). Results are from three independent experiments. *, p<0.05, ***, p<0.001. **G)** Images showing cell morphology of beige differentiated SVFs isolated from male and female iWATs. **H)** Body weight and lean mass of male and female mice before (RT) and after 24 hr cold exposure (COLD). N=6,6. **, p<0.01. **I-K)** Individual mice data for oxygen consumption (VO2 ml/hr), carbon dioxide production (VCO2 ml/hr), and energy expenditure (EE kcal/hr) (I), respiratory exchange ratio (RER) and generalized linear model (GLM)-based regression plots of RER with total body weight (Total), lean mass (Lean) and fat mass (Fat) as co-variates (ANCOVA) showing p-value for Mass effect, Group effect, and Interaction effect. *, p<0.05 (J), and total food consumed (kcal), locomotor activity (beam breaks), and total distance in cage (m) (K) of male and female mice exposed to 22°C for 21 hr and 4°C for 24 hr in Sable Promethion metabolic chambers (12 hr light/dark cycle, 45 hr total duration, white bar represents light cycle and grey bar represents night cycle). N=6,6.

**Extended Data Figure 4.**
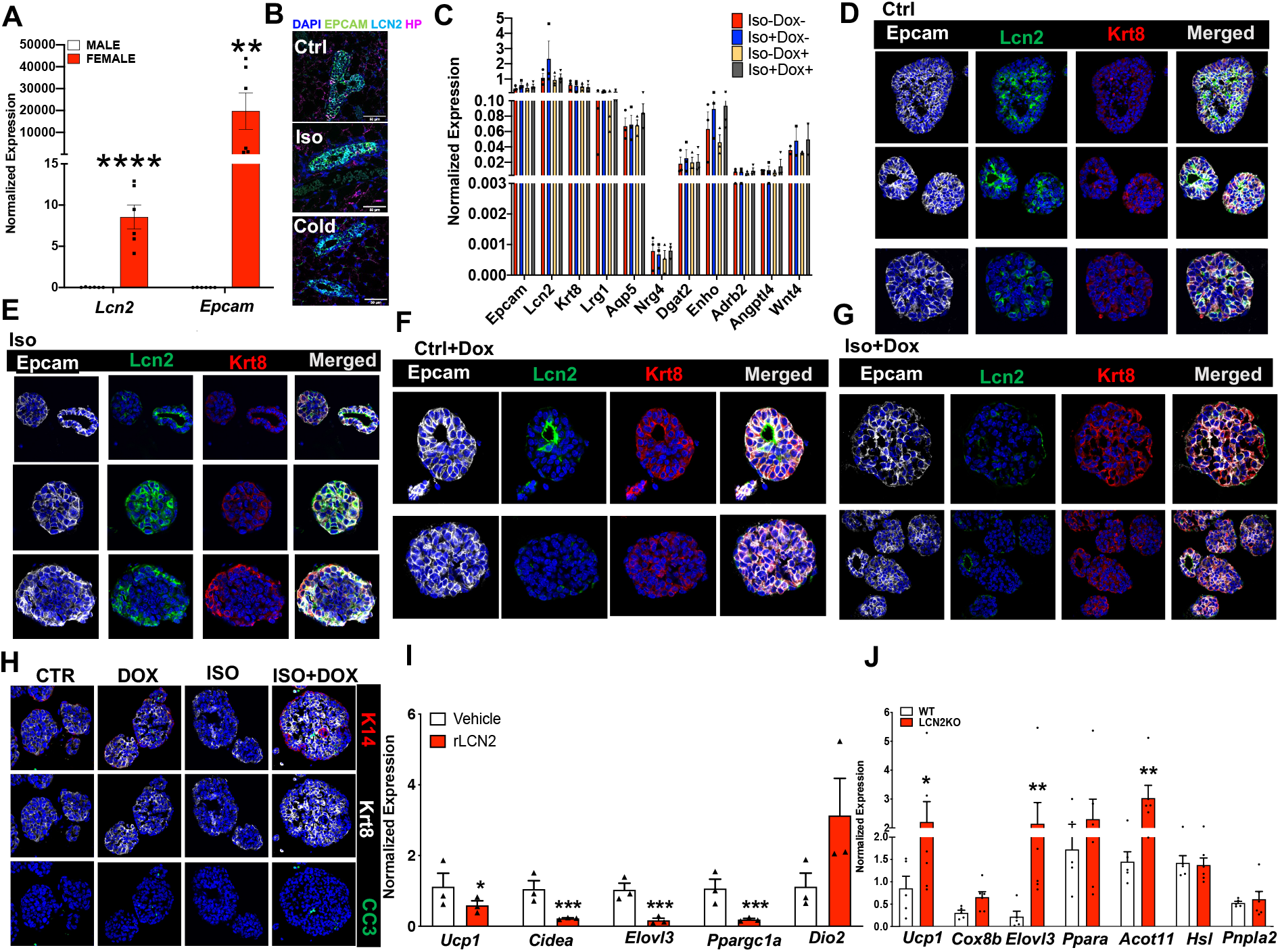
LCN2 is a cold-induced mammokine involved in blocking cold-induced adipocyte UCP1 expression. **A)** Real-time qPCR of indicated genes from 24 hr cold-exposed male and female iWATs. **, p<0.01; ***, p<0.001. **B)** RNAScope FISH (see Materials and Methods) of indicated probes from mgWAT of RT or 24 hr isoproterenol or cold-exposed mice. **C)** Real-time qPCR of indicated genes in organoids derived from K8rtTA-DTA mice treated with and without ISO and Doxycycline (Dox). Results from 3 organoid experiments. **D-H)** Confocal images of immunostaining of indicated antibodies in organoids derived from K8rtTA-DTA mice treated with and without Iso and Dox. Representative images from 2-3 organoid experiments. **I)** Real-time qPCR of indicated genes from beige differentiation mgWAT SVFs derived from *Lcn2* KO mice treated with and without recombinant Lcn2 (rLcn2). *, p<0.05; ***, p<0.001 **J)** Real-time qPCR of indicated genes from 24 hr cold exposed WT and *Lcn2* KO mgWATs. N=5,5. *, p<0.05; **, p<0.01

