## Supplementary Movie 1 for "Mammary ductal epithelium controls cold-induced adipocyte thermogenesis"

### Slide 1
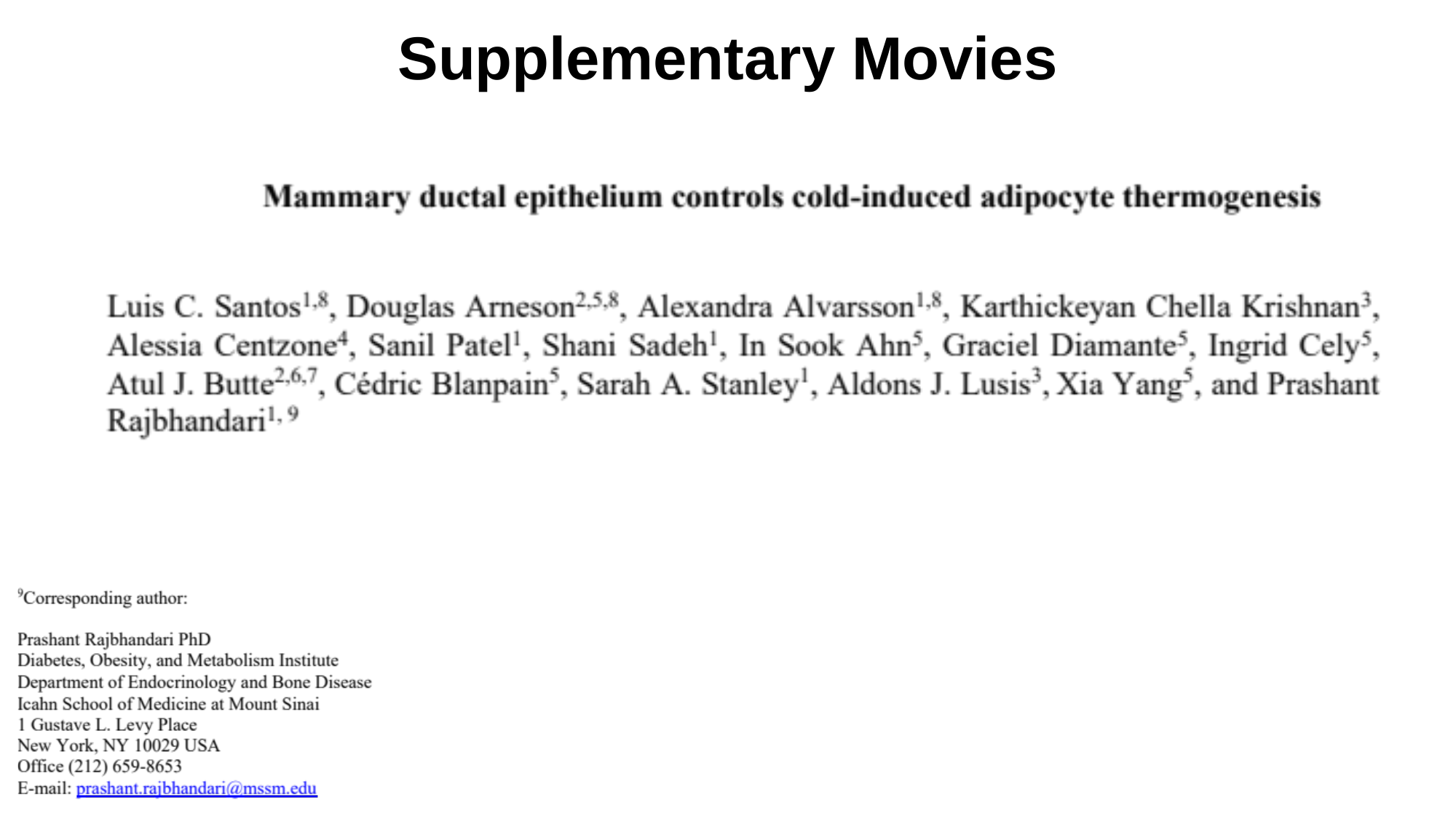

Supplementary Movies

### Slide 2
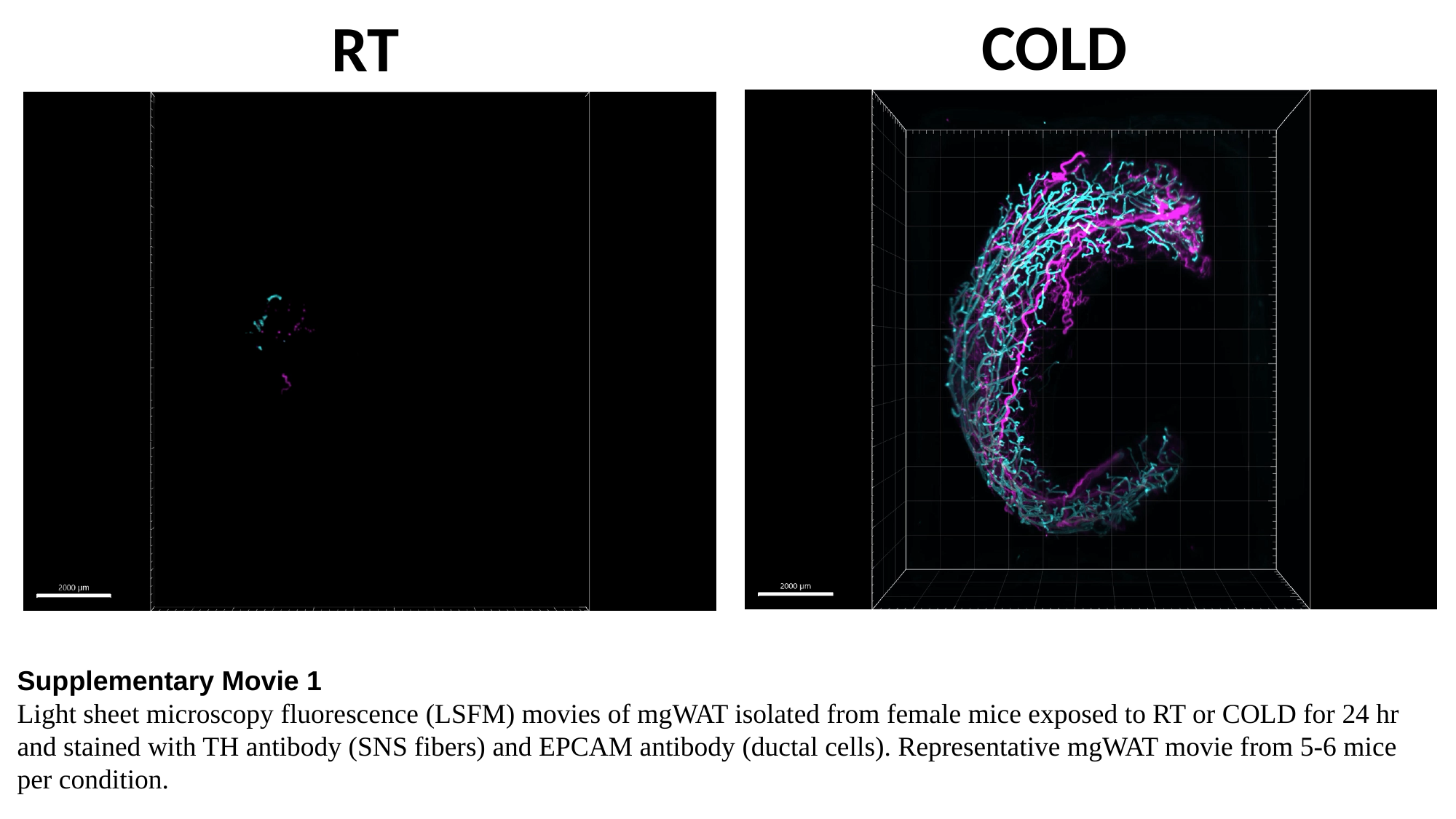

COLD
RT
Supplementary Movie 1
Light sheet microscopy fluorescence (LSFM) movies of mgWAT isolated from female mice exposed to RT or COLD for 24 hr and stained with TH antibody (SNS fibers) and EPCAM antibody (ductal cells). Representative mgWAT movie from 5-6 mice per condition.
Supplemental Movie 2
