## Supplementary Movie 2 for "Mammary ductal epithelium controls cold-induced adipocyte thermogenesis"

### Slide 1
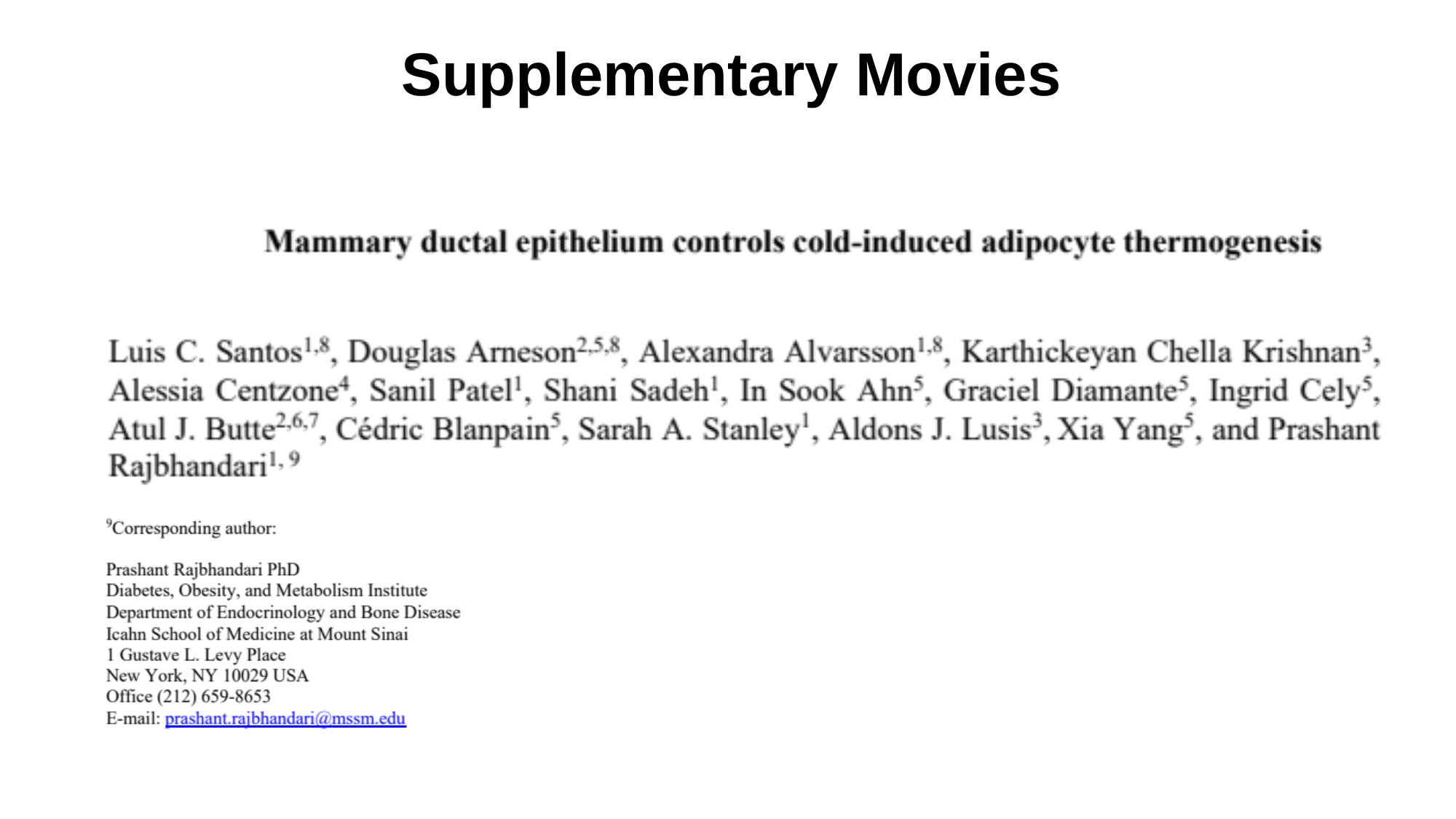

Supplementary Movies

### Slide 2
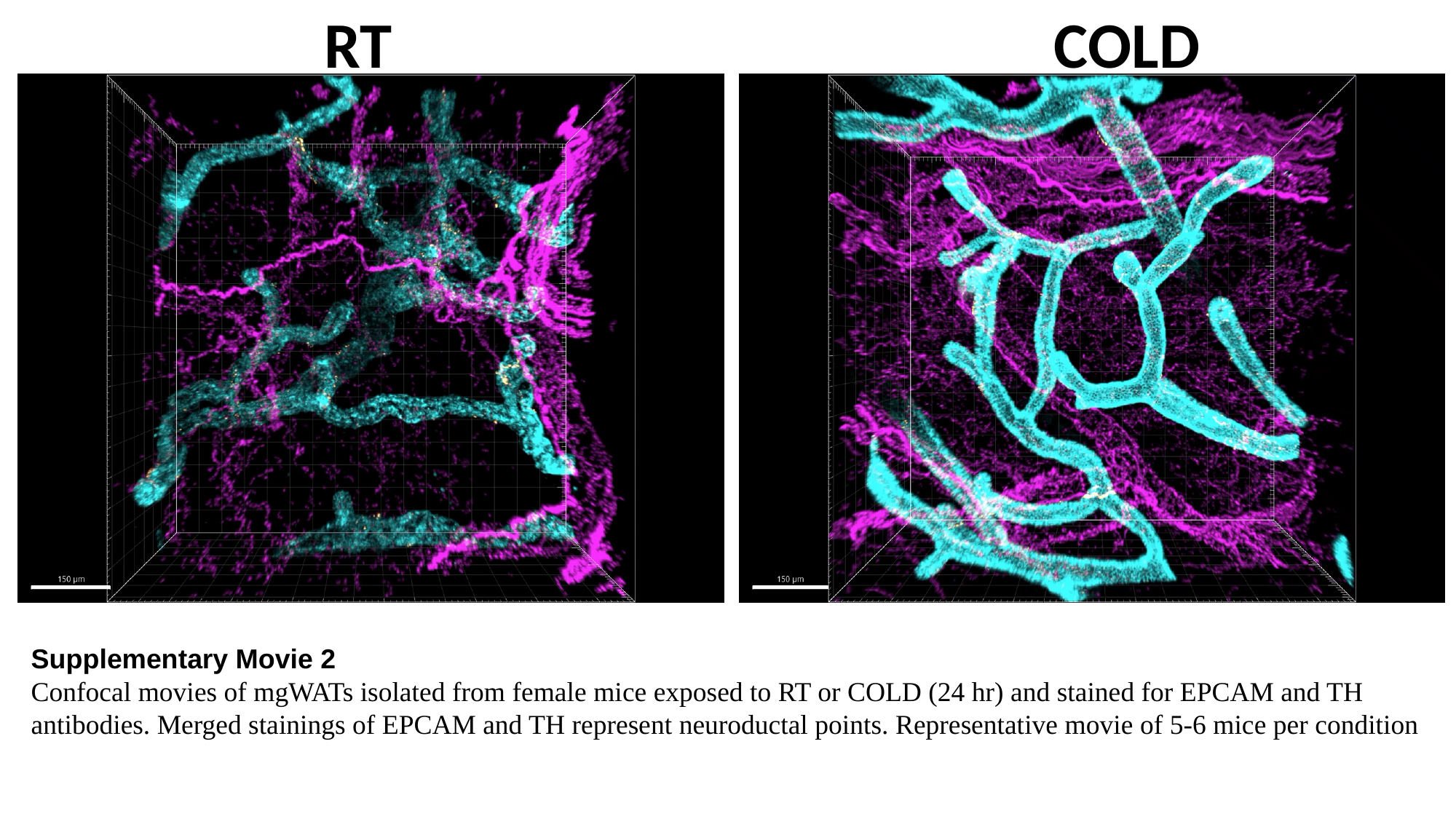

RT
COLD
Supplementary Movie 2
Confocal movies of mgWATs isolated from female mice exposed to RT or COLD (24 hr) and stained for EPCAM and TH antibodies. Merged stainings of EPCAM and TH represent neuroductal points. Representative movie of 5-6 mice per condition
Supplemental Movie 2
